# Connectivity patterns of task-specific brain networks allow individual prediction of cognitive symptom dimension of schizophrenia and link to molecular architecture

**DOI:** 10.1101/2020.07.02.185124

**Authors:** Ji Chen, Veronika I. Müller, Juergen Dukart, Felix Hoffstaedter, Justin T. Baker, Avram J. Holmes, Deniz Vatansever, Thomas Nickl-Jockschat, Xiaojin Liu, Birgit Derntl, Lydia Kogler, Renaud Jardri, Oliver Gruber, André Aleman, Iris E. Sommer, Simon B. Eickhoff, Kaustubh R. Patil

**Author notes:** Correspondence should be addressed to: Simon B. Eickhoff, Institute of Systems Neuroscience, Heinrich Heine University Düsseldorf, 40225 Düsseldorf, Germany & Institute of Neuroscience and Medicine, Brain and Behaviour (INM-7), Research Center Jülich, 52428 Jülich, Germany. Or Ji Chen Institute of Systems Neuroscience, Heinrich Heine University Düsseldorf, 40225 Düsseldorf, Germany & Institute of Neuroscience and Medicine, Brain and Behaviour (INM-7), Research Center Jülich, 52428 Jülich, Germany.

## Abstract

**Background:** Despite the marked inter-individual variability in the clinical presentation of schizophrenia, it remains unclear the extent to which individual dimensions of psychopathology may be reflected in variability across the collective set of functional brain connections. Here, we address this question using network-based predictive modeling of individual psychopathology along four data-driven symptom dimensions. Follow-up analyses assess the molecular underpinnings of predictive networks by relating them to neurotransmitter-receptor distribution patterns.

**Methods:** We investigated resting-state fMRI data from 147 schizophrenia patients recruited at seven sites. Individual expression along negative, positive, affective, and cognitive symptom dimensions was predicted using relevance vector machine based on functional connectivity within 17 meta-analytic task-networks following a repeated 10-fold cross-validation and leave-one-site-out analyses. Results were validated in an independent sample. Networks robustly predicting individual symptom dimensions were spatially correlated with density maps of nine receptors/transporters from prior molecular imaging in healthy populations.

**Results:** Ten-fold and leave-one-site-out analyses revealed five predictive network-symptom associations. Connectivity within theory-of-mind, cognitive reappraisal, and mirror neuron networks predicted negative, positive, and affective symptom dimensions, respectively. Cognitive dimension was predicted by theory-of-mind and socio-affective-default networks. Importantly, these predictions generalized to the independent sample. Intriguingly, these two networks were positively associated with D_1_ dopamine receptor and serotonin reuptake transporter densities as well as dopamine-synthesis-capacity.

**Conclusions:** We revealed a robust association between intrinsic functional connectivity within networks for socio-affective processes and the cognitive dimension of psychopathology. By investigating the molecular architecture, the present work links dopaminergic and serotonergic systems with the functional topography of brain networks underlying cognitive symptoms in schizophrenia.

## Introduction

Precise conceptualization of schizophrenia symptoms in terms of their underlying dimensional structure and associated neurobiology remains a challenge (1), as prior work focused on to subscales (positive, negative, and general symptoms) of the Positive and Negative Syndrome Scale (PANSS) (2) has not provided a clear understanding of underlying brain circuitry (3-5). We recently introduced a novel four-dimensional conceptualization of schizophrenia symptomatology which is stable and generalizable across populations and settings (5). As each symptom dimension captures a different clinical facet of schizophrenia (6-8), we would expect these to show differential relationships with functional brain networks. Identification of robust symptom-brain relationships (e.g., connectivity patterns and molecular substrates) is important for the development of reliable biomarkers for targeted treatments of different symptom dimensions. Previous studies proposed that abnormal brain connectivity might be a precipitating factor for schizophrenia symptoms (9,10), questioning region-based analyses but resonating with the dysconnection hypothesis (9-12).

Although resting-state functional MRI (fMRI) reveals broad patterns of impaired brain function that may underlie the pathophysiology of schizophrenia (12-15), the link between targeted symptom dimensions and associated connectivity patterns within distinct functional systems remain largely unknown. Pioneering work has explored symptom-brain associations based on regional activity and intrinsic connectivity networks (ICNs) using univariate group-level correlative approaches, but the results have been largely inconsistent (16-19). The clinical complexity of schizophrenia together with the differences in patient populations, study settings, scanners and scanning protocols across sites may have led to divergent results, posing a major challenge for establishing generalizable network-symptom relationships. Application of multivariable machine-learning and cross-validation strategies to multi-site data and validation of the resulting models on independent datasets is thus needed to derive robust network-symptom associations (20).

It needs to be cautioned, though, that ICNs cannot readily be interpreted relative to cognitive and mental processes due to their unconstrained and task-independent nature (21). In contrast, meta-analytic functional networks are derived from task-activation data, i.e., the identified networks consist of brain areas robustly engaged in specific tasks and therefore mental processes (22,23). Meta-analytic networks thus provide a promising avenue to characterize association between functionally meaningful systems and specific symptom dimensions. Considering these advantages, we here combined multivariable machine-learning and meta-analytically defined networks to explore predictive relationships between network-specific connectivity patterns and individual expressions of different dimensions of psychopathology.

Functional brain systems are known to relate with molecular architecture (24-27). In order to facilitate a link to treatment, we also explored whether robustly symptom-related functional networks would in turn relate to the spatial topography of underlying molecular features. Specifically, connectivity-neurotransmitter coupling has been proposed and observed in healthy populations (28,29). Similarly, network dysconnectivity in schizophrenia has been associated with altered neurotransmission (30,31) with several pathways involving dopaminergic, serotonergic, gamma-aminobutyric acid(GABA)-ergic, and glutamatergic systems reported to be compromised (32-35). Here, it is interesting to note that current anti-psychotic drugs primarily targeting the dopamine system are primarily effective against positive but less so for negative or cognitive symptoms (36,37). Understanding the molecular substrate of specific dimensions of psychopathology may thus provide leads on new treatment strategies.

We therefore assessed a broad range of meta-analytic networks relating to social, affective, executive, memory, language, and sensory-motor functions with respect to their predictive power for individual positive, negative, affective and cognitive symptom-dimensions in schizophrenia. Machine-learning approaches with a stringent validation sequence of 10-fold cross-validation, leave-one-site-out analyses, and generalization to an independent sample was implemented to identify robust association patterns between the probed networks and the four dimensions of psychopathology (6). Subsequently, whole-brain density maps of nine receptors/transporters from prior in vivo molecular imaging studies were employed to investigate the molecular architecture spatially coupled to the identified predictive networks.

## Materials and Methods

### Sample

A total of 147 schizophrenia patients from seven centers located in Europe (Aachen [Aachen-1 and Aachen-2], Göttingen, Groningen, Lille, and Utrecht) and the USA (COBRE, Albuquerque, NM) represented the main sample (Table S1; Supplement). These sites differed significantly in illness duration (*p*<0.001)(Table S1). An independent sample with 117 schizophrenia patients (Table S2; Supplement) retrieved from the Bipolar-Schizophrenia Network on Intermediate Phenotypes (B-SNIP) database (38) was used for independent validation of the predictive models. For both samples, diagnosis of schizophrenia was established based on the DSM-IV, the DSM-IV-TR, or the ICD-10 criteria (see Supplement and [6]). These international datasets cover a broad range of clinical states, settings, and medical systems, facilitating identification of robust network-symptom associations. Current drug dosages of antipsychotic medication were olanzapine-equivalent transformed (39). For each site, subjects gave written informed consent and study approval was given by the respective ethics committees/institutional review boards. Additional approval to pool and re-analyze data was provided by the ethics committee of the University of Düsseldorf, Germany.

### Calculation of dimensional symptom scores

Severity of psychopathology was assessed using the PANSS (2). The 30 PANSS items were compressed into four (negative, positive, affective, and cognitive) symptom dimensions (Figure S1A) identified in our prior factorization analysis on two large, multi-site schizophrenia samples as stable and well-generalizable across populations, settings, and medical systems (6). The original item-by-subject matrix was projected onto this dimensional-structure of PANSS to yield the dimensional symptom scores for each subject (as implemented in *DCTS:* http://webtools.inm7.de/sczDCTS/). Higher scores on a dimension indicate more-severe symptoms (Figure S1B).

### Definition of functional brain networks

Seventeen functional networks, which cover a broad range of domains reflecting cognitive, socio-affective, and sensory-motor functions that have been implicated in schizophrenia, were employed (Table S3). These networks were based on coordinate-based meta-analyses (21,22) and represent regions demonstrating convergent activations associated with specific functional domains across many prior task-fMRI studies. They hence provide the best ―a *priori*‖ estimate of the location of specific functional networks and hence here assessed by resting-state fMRI in new subjects. For convenience, we grouped these 17 networks into six broad functional domains (Figure 1&Table S3), though it must be stressed that each network was analyzed separately.

**Figure 1.**
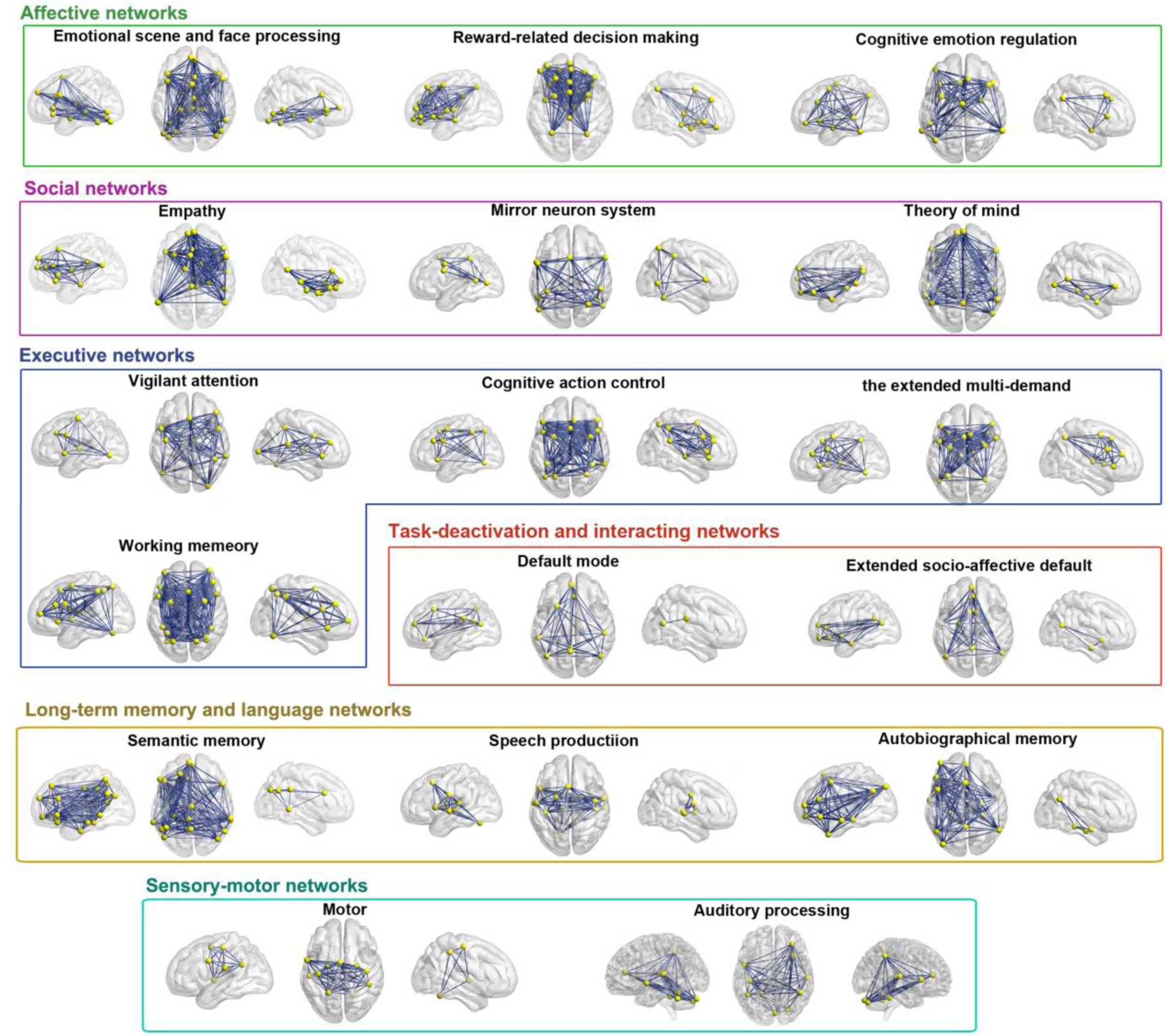
Overview of the 17 functional networks derived from prior coordinate-based meta-analytical studies. Networks were assigned to six broad domains according to their main functional roles, as implicated in the tasks included in the source publications of these meta-analytical networks, in multiple neuro-cognitive and socio-affective processes. We also note that networks such as the task-deactivation default-mode (DMN) network (22) and the extended socio-affective-default (eSAD) network which is derived from DMN regions (71) can be engaged during multiple processes (69,70). Details can be found in Supplementary Table S3. Yellow nodes represent the spheres created from the coordinates with 6mm radius and blue lines denote the pair-wise connections between the nodes. Connections within each network were used separately in our predictive modeling to investigate network-specific relationships with individual dimensions of psychopathology.

**Figure 2.**
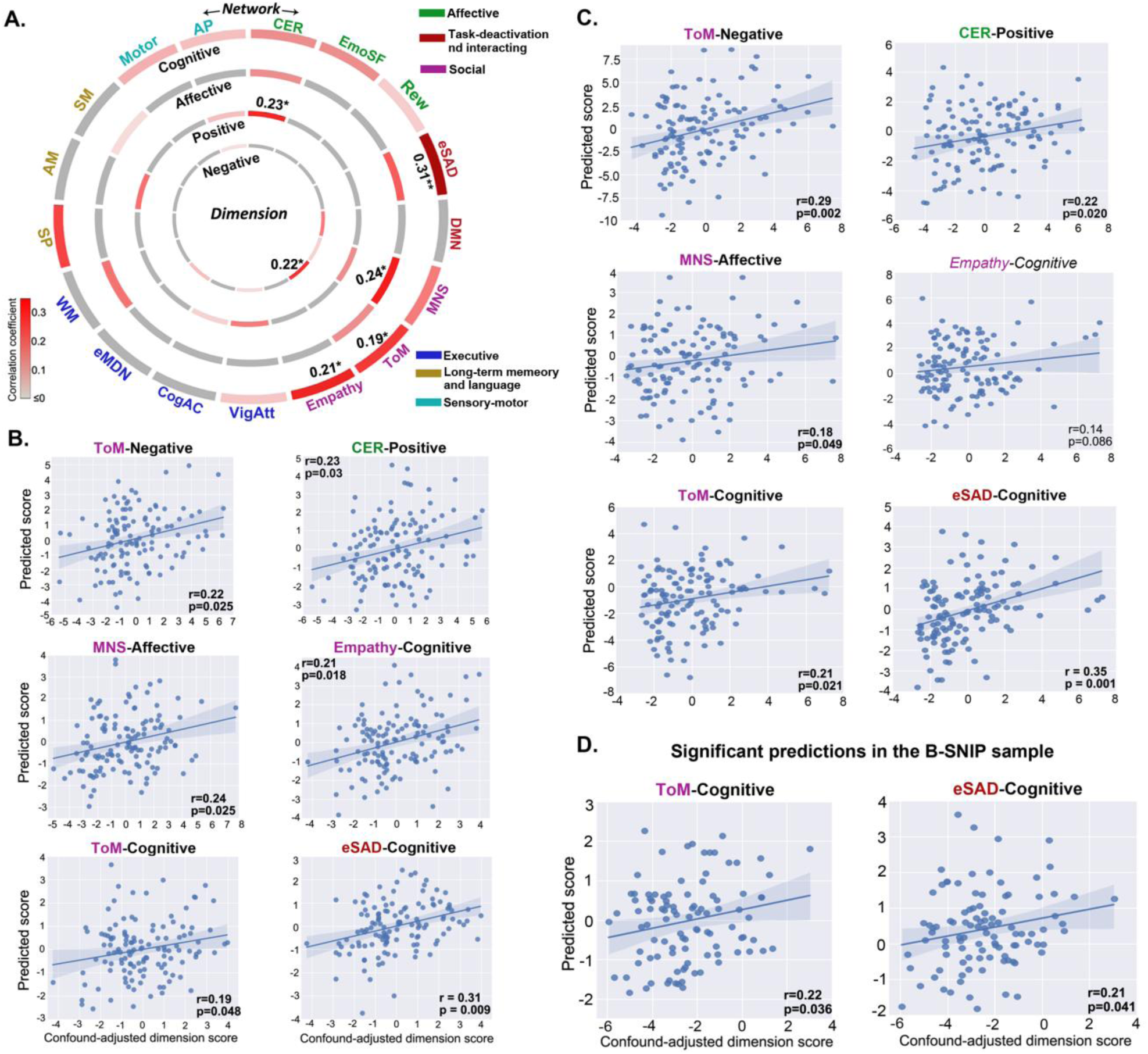
Multivariable prediction of the four symptom dimensions from within-network resting-state functional connectivity using relevant vector machine. **A)** Circos plot shows the 500x repeated 10-fold cross-validation results for the main sample. Correlations between the actual (confound-adjusted) dimensional symptom scores and their predictions are color-coded from light grey (0) to dark red (0.35), *\*p*<*0*.*05, **p*<*0*.*01*. **B)** Scatter plots show the six significant predictions identified by 10-fold cross-validation in main sample. **C)** Scatter plots show the leave-one-site-out cross-validation results for the six significant predictions identified by 10-fold cross-validation in main sample. Apart from the prediction of cognitive dimension from the rs-FC within the empathy network, other predictions were all confirmed by leave-one-site-out with significant correlations identified. **D)** Scatter plots show the significant predictions in the independent B-SNIP sample. Models trained within the main sample were used for this validation analysis in B-SNIP. Shaded areas represent 95% confidence intervals. Abbreviations: EmoSF, emotional scene and face processing; Rew, reward-related decision making; CER, cognitive emotion regulation; ToM, theory-of-mind; MNS, minor neuron system; DMN, default mode network; eSAD, extended socio-affective default; VigAtt, vigilant attention; CogAC, cognitive action control; eMDN, the extended multi-demand networks; SM, semantic memory; SP, speech production; WM, working memory; AM, autobiographical memory; APN, auditory processing network.

### FMRI data preprocessing

All resting-state fMRI scans (Tables S4&S5) were preprocessed using SPM12 (http://www.fil.ion.ucl.ac.uk/spm) (see [6] and Supplement). After excluding subjects with excessive head-motion (40) or poor image quality (details in Supplement), 126 and 100 schizophrenia patients were retained in the main and the B-SNIP validation sample, respectively (Table 1). Head motion differed significantly between the sites in the main sample (*p*=0.003; one-way ANOVA) and between the main and the B-SNIP samples (*p*<0.001; two-sample t-test) but did not correlate with the residuals of any symptom dimensions after adjusting for age/gender/site. Still, we adjusted head motion effects in predictive modeling as a conservative approach to rule out (any) possible predictability of symptom dimensions due to movement.

**Table 1.** Demographic and clinical characteristics of schizophrenia patients used for predictive analysis.

| **Characteristics** | **Main sample**<br>**(N=126, seven**<br>**sites)** | **The independent B-SNIP**<br>**sample for validation**<br>**(N=100, two sites)** | **p-value** |
| --- | --- | --- | --- |
| **Demographic** |  |  |  |
| Age (years) | 34.19 (11.45) | 34.28 (12.31) | 0.948 |
| Gender (male/female) | 92/34 | 71/29 | 0.767 |
| Illness during (years) | 10.48 (9.87) | 12.35 (11.13) | 0.187 |
| **PANSS subscales** |  |  |  |
| Positive | 14.86 (5.35) | 14.91 (5.72) | 0.945 |
| Negative | 14.76 (5.85) | 15.01 (5.64) | 0.743 |
| General | 30.25 (10.13) | 27.98 (7.46) | 0.062 |
| Illness severity (total<br>PANSS) | 59.87 (18.11) | 57.90 (15.78) | 0.390 |
| **Scores on the dimensions**<br>**of PANSS** |  |  |  |
| Negative | 2.76 (2.44) | 2.81 (2.23) | 0.859 |
| Positive | 3.26 (2.36) | 3.28 (2.55) | 0.954 |
| Affective | 3.33 (2.33) | 2.69 (1.71) | **0.022** |
| Cognitive | 2.49 (1.92) | 2.72 (1.79) | 0.357 |
| **Medication** |  |  |  |
| Atypical antipsychotics | 97 (75.8%) | 71 (71.0%) |  |
| Typical antipsychotics | 5 (3.9%) | 4 (4.0%) |  |
| Both A & T | 7 (5.5%) | 13 (13.0%) |  |
| None or unknown | 19 (14.8%) | 12 (12.0%) |  |
| Current antipsychotic<br>medication<sup>e</sup> | 19.23 (11.91) | 18.96 (13.47) |  |

Data are mean (SD), or n (%). *p* Values in bold indicate a significance of $p < 0.05$ . Except for gender, which was based on chi-square test, other statistics were all based on two-sample *t* test. Of note, because the detailed medication information was missing for several patients in different proportions for the three datasets, statistical comparisons were not conducted.
PANSS, Positive and Negative Syndrome Scale;
<sup>e</sup>Demonstrated in olanzapine-equivalent dosage (mg/day).

White matter and CSF signals as well as 24 head-motion parameters (41,42) were regressed out from the overall fMRI time-series (42) but the global mean signals were not removed given ongoing controversies (43,44). The first eigenvariate of the time-series from all the voxels within a 6-mm sphere around each node was extracted (45,46). For each network, the between-node resting-state functional connectivity (rs-FC) was then calculated as Fisher’s z-transformed Pearson’s correlation between the eigenvariates.

### Prediction of symptom dimensions using network rs-FC

Multivariable regression via a relevance vector machine (RVM)(47) was implemented and evaluated using 500x repeated 10-fold cross-validation, individually for each combination of functional network and symptom dimension in the main sample. That is, we assessed the capability of each network’s rs-FC to predict each of the four symptom dimensions in held-out patients. Importantly, RVM is a sparse learning method, i.e., only a few of the feature weights learned by RVM are non-zero, lending interpretability as to which features (connections) are predictive. In keeping with the recommended strategy (48), both the symptom dimensional-scores and rs-FC features were adjusted for confounding effects of age, gender, site, and head-motion (DVARS). To avoid data-leakage within cross-validation (49), confound regression models were learned only on the training-set and then applied on both training and test data (50,51). The RVM model was then trained on confound-adjusted training data and applied to the confound-adjusted held-out test data. The folds were stratified to accommodate different sample sizes across sites. Prediction performance was evaluated using *Pearson’s* correlation between the (adjusted) scores and their predictions. Significantly predictive associations were further validated for their generalizability across sites using leave-one-site-out cross-validation following the same schematic but training on all sites but one and testing on the left-out site (cf. Supplement). Statistical significance of the cross-validation-based correlations was determined through 1000 permutations by shuffling the symptom dimensional-scores (lowest *p*=0.001, right-tailed; Supplement) (49,52).

Critically, we validated the associations confirmed as predictive in the leave-one-site-out cross-validation in the independent B-SNIP sample. For this, we trained RVM models on the significantly predictive networks in the entire main sample and then, without further fitting or modification, applied them to held-out B-SNIP data. Robust associations passing this strict three-step validation procedure were then further assessed as described below.

In addition, we performed control analyses by repeating all validation procedures by including illness duration or olanzapine-equivalent dosage as confounds. For comparison, we also predicted the original three PANSS subscales (positive, negative, and general psychopathology) using the same three-step validation procedure.

### Identification of reliably predictive connections and subnetworks

The intrinsic feature selection in RVM through its sparse modeling was leveraged to identify reliably predictive connections and the potential subnetworks formed by them. A connection was identified as reliably predictive when it had non-zero weights in: i) at least 80% of the 10-fold cross-validation repetitions, ii) at least six out of the seven (i.e., >80%) leave-one-site-out analyses, and iii) the models trained on the entire main sample for validation in B-SNIP. The cutoff of 80% is suggested as a conservative threshold to select most relevant variables in both real and simulated data (53). To assess the predictive capacity of the subnetworks, RVM models were trained using the rs-FC of the subnetworks on the main sample and tested in B-SNIP.

### Spatial correlation with receptor/transporter density estimates

Finally, we evaluated the topographical relationship between network-node location and the distribution of several receptor/transporter systems, assessing if any receptors/transporters were highly-expressed in the identified networks relative to the entire brain. This was tested by comparing the average receptor/transporter density across all nodes within a given network against a null-distribution based on 1000 random network configurations generated by re-distributing the nodes throughout the grey matter while preserving the between-node distances (±6mm tolerance). Seven dopamine and serotonin receptors (dopaminergic: D_1_ and D_2/3_; serotonergic: 5-HT1a, 5-HT1b, and 5-HT2a) and transporters (dopamine transporter and 5-HTT serotonin reuptake transporter), together with F-DOPA (a reflection of presynaptic dopamine-synthesis-capacity) and the GABAergic receptor GABAa were investigated. These three neurotransmitter systems have all been implicated in schizophrenia (32,34,35), while here we tested for more specifically the receptors/transporters. Density estimates for these receptors/transporters were derived from average group maps of healthy volunteers scanned in prior multi-tracer molecular imaging studies (Supplement). For comparability, these maps, in MNI152 space, were all resampled to an isotropic 2mm spatial resolution as in our fMRI data and linearly rescaled to a minimum of 0 and a maximum of 100.

Furthermore, the significantly higher expressed receptors/transporters were entered into a spatial correlation analysis (54,55) calculated as rank correlation between the node importance scores and receptor/transporter densities calculated for these nodes (Figure 4B). The node importance score was calculated by summing the selection frequency of the connections of each node derived from the repeated 10-fold cross-validation. Bootstrap analysis was conducted to ensure robustness. To establish the statistical significance of a spatial correlation against chance, spatial permutation test was employed where the null distribution was estimated based on the correlations between the node importance scores for a given network and the nodal receptor/transporter densities extracted from1000 simulated (random) networks (Supplement).

**Figure 3.**
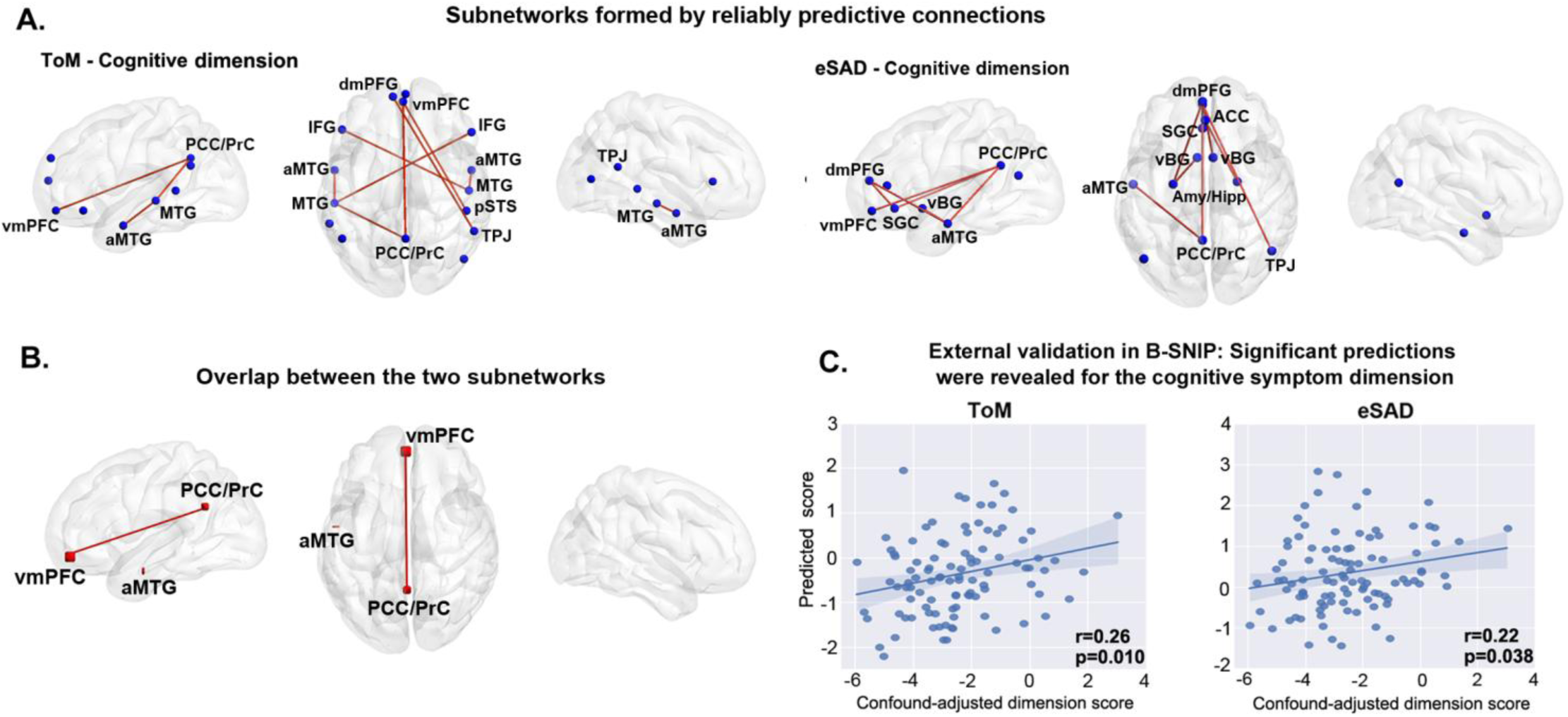
Reliably predictive connections for the two networks that robustly predicted the cognitive dimension and validation of the formed subnetworks in the independent sample. **A)** Reliably predictive connections selected in more than 80% of the different cross-validation runs (repeated 10-fold and leave-one-site out) and in the final models trained on the entire main sample for validation in B-SNIP. **B)** Overlapping between the two subnetworks. **C)** Scatter plots show the significant correlations in B-SNIP using the models trained within the main sample. Abbreviations: ToM, theory-of-mind; eSAD, extended socio-affective default; CER, cognitive emotion regulation, MNS, mirror neuron system. Amy, amygdala; Hipp, hippocampus; vmPFC, ventro-medial prefrontal cortex; dmPFC, dorso-medial prefrontal cortex; mFG, medial frontal cortex; aMTG, anterior middle temporal gyrus, IFG, inferior frontal gyrus; TPJ, temporo-parietal junction, PCC, posterior cingulated cortex, PrC, precuneus; SGC, subgenual cingulate cortex, vBG, ventral basal ganglia; ACC, anterior cingulated cortex.

**Figure 4.**
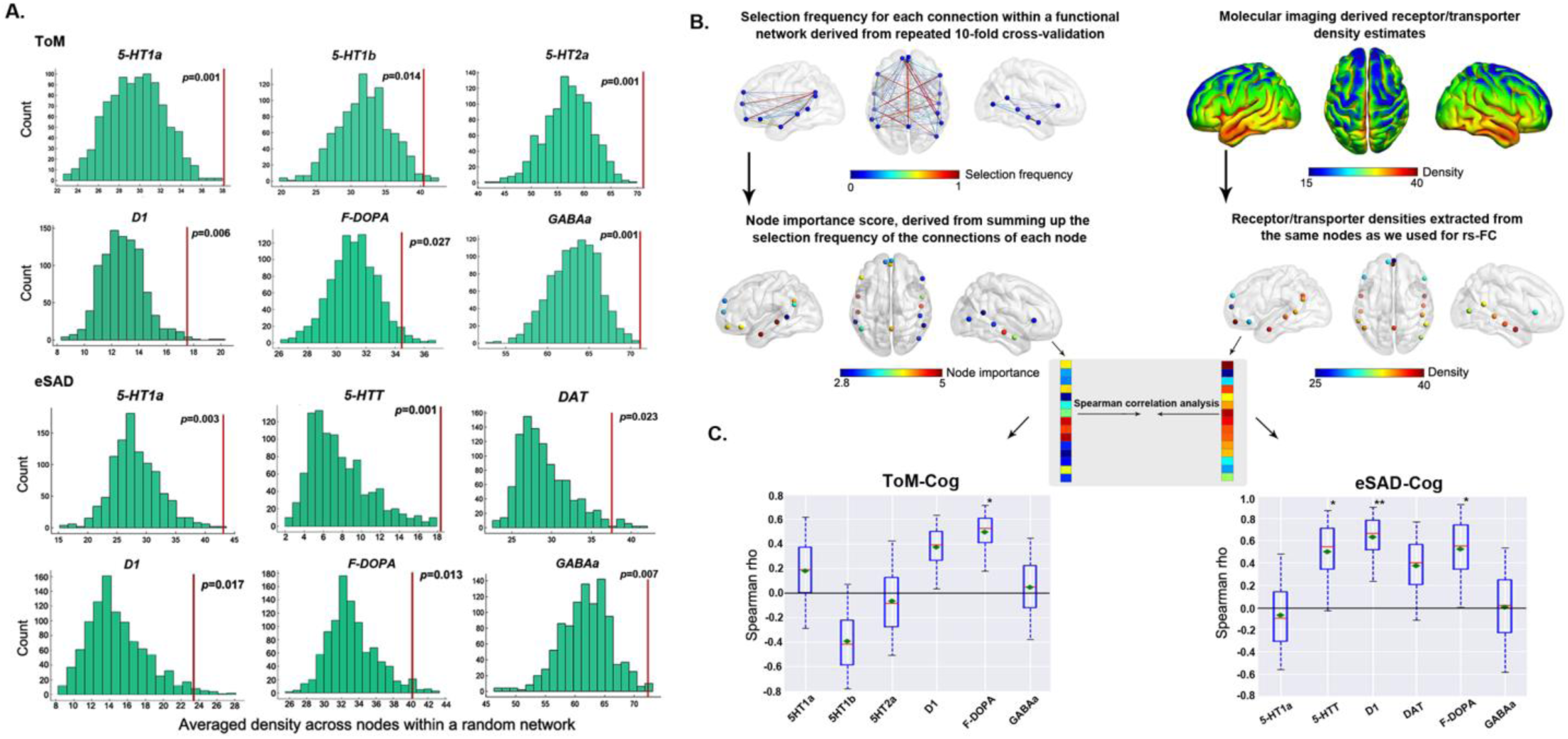
Significantly highly-expressed receptors and transporters at the nodes of the robustly predictive networks relative to the entire brain, as well as the schematic and results for the spatial correlation analysis with receptor/transporter density maps. **A)** Histograms show the null distributions for the receptor/transporter densities of the 1000 simulated (random) networks. Red lines indicate the true averaged receptor/transporter densities across the different nodes within real networks. **B)** Procedure for conducting the spatial correlation analysis between network nodes and receptor/transporter density maps. **C)** Bootstrapped Spearman correlations (repeated 10,000 times) between the node importance score and the nodal receptor/transporter density estimates for the two identified networks which robustly predicted the cognitive symptom dimension. Bootstrap nodes were drawn with replacement from the real networks, and then the correlation analysis was done on them. Boxes refer to the Spearman rho values. The red line depicts the median, the green diamond depicts the mean, and the whiskers represent the 5th and 95th percentiles. Significant correlations derived from spatial-level permutation tests are marked with an asterisk (\**p*<0.05, \*\**p*<0.01). Abbreviations: ToM, theory-of-mind; eSAD, extended socio-affective default; Cog, cognitive dimension.

## Results

### Network-based prediction of specific symptom dimensions

Predictive modeling with a stringent three-step validation revealed specific networks associated with the four dimensions of psychopathology. In the first step (10-fold cross-validation), the negative, positive, and affective symptom dimensions could be significantly predicted using the rs-FC within the theory-of-mind (ToM), the cognitive emotion regulation (CER), and the mirror neuron networks, respectively, while the cognitive dimension was predicted by three networks, ToM, empathy and eSAD (Figure 2A, B). These six networks-symptom associations (Figure 2B) were then tested in the second validation step, i.e., leave-one-site-out. Except for the empathy-cognitive prediction, all predictive associations were validated by significant correlations between the observed (confound-adjusted) scores for the individual symptom-dimensions and their predictions (Figure 2C).

As the 10-fold cross-validation and leave-one-site-out analyses were both performed in the main sample, they may still be optimistic with respect to the generalization to new patient populations. Therefore we added a third validation step for the five leave-one-site-out validated associations using a completely independent sample. Three of the five leave-one-site-out validated associations (ToM-negative, CER-positive, MNS-affective) were not confirmed in this step. Training RVM models on the entire main sample and testing it in the B-SNIP dataset revealed that the ToM and eSAD networks were significantly predictive of cognitive symptoms (Figure 2D).

In complementary analyses, no significant effects of illness duration or olanzapine-equivalent dosage on the four symptom-dimension scores were found (all *p*>0.05 in the fitted general linear models with additional covariates of age, gender, site, and head motion). Conversely, controlling for illness duration or olanzapine-equivalent dosage did not alter the overall predictive patterns. Highlighting the utility of our four-dimensional conceptualization of schizophrenia symptomatology (6), when considering the traditional three PANSS subscales, 10-fold cross-validation and leave-one-site-out experiments on the main sample only revealed two predictive patterns: the ToM network predicted the negative subscale, and the eSAD network predicted the general psychopathology subscale (Figure S2A, B). However, these two associations were not generalizable to B-SNIP (Figure S2C).

### Reliably predictive connections and the predictiveness of subnetworks

Ten connections within eSAD and eight connections within ToM were identified as consistently relevant for predicting the cognitive symptoms (Figure 3A; Table S6), i.e., were selected in more than 80% of the different cross-validation runs (repeated 10-fold and leave-one-site out) and in the final models trained on the entire main sample for validation in B-SNIP. The ensuing ToM and eSAD subnetworks featured three spatially overlapping nodes located in the ventro-medial prefrontal cortex (vmPFC), left middle temporal gyrus (MTG), and posterior cingulate cortex (PCC)/precuneus and highlighted the vmPFC-PCC/precuneus connection (Figure 3B; Table S7). In turn, connections to subcortical regions including bilateral amygdala/hippocampus and the ventral striatum were specific to the eSAD subnetwork.

A model based on the identified eSAD subnetwork showed almost identical prediction performance for the B-SNIP data compared to the aforementioned one trained on whole eSAD network (Figure 3C, note that subnetwork definition was only based on the main sample, i.e., there is no leakage of information about the B-SNIP data). Interestingly, compared to whole ToM network (Figure 2D), the ToM subnetwork demonstrated an improved performance (Figure 3C) in the prediction of the cognitive dimension in B-SNIP. This confirms the power of sparse modeling in RVM to identify the truly relevant features.

### Relationship to molecular architecture

Results showed that eight receptors/transporters (Figure 4A) are highly expressed in both, or in either of the ToM and eSAD networks than in the simulated networks with perturbed spatial configuration of nodes (histograms for within- and between-network distance of nodes shown in Figure S3). Spatial correlation between the node importance (Table S8) and the spatially corresponding density of those significant receptors/transporters revealed a relationship to both the dopaminergic and the serotonergic systems (Figure 4C). Specifically, the nodes of ToM that showed higher importance in predicting the cognitive dimension tracked with higher dopamine-synthesis-capacity (*r*=0.54, *p*=0.02). The prediction importance of the nodes within the eSAD network correlated positively with densities of D_1_ (*r*=0.66, *p*=0.007) and 5-HTT (*r*=0.53, *p*=0.046), as well as dopamine-synthesis-capacity (*r*=0.54, *p*=0.036). The significance was corroborated by bootstrapped confidence intervals.

## Discussion

By employing predictive modeling to multi-site fMRI data with a strict sequence of out-of-sample validation steps (repeated 10-fold, leave-one-site-out, independent dataset), two a priori, meta-analytically defined functional networks, theory-of-mind (ToM) and extended socio-affective default (eSAD), were identified as significantly and robustly predictive of the cognitive symptom dimension. In contrast, prediction using the original PANSS subscales failed to generalize to the independent sample, supporting the notion that these traditional dimensions do not correspond well to underlying neurobiology. Through the implicit feature selection of RVM, reliably predictive connections were identified which constituted subnetworks connecting nodes mainly distributed in the (medial) prefrontal cortex, PCC, temporal regions as well as subcortical structures. Moreover, higher densities of D_1_ dopamine receptor and 5-HTT serotonin transporter as well as elevated dopamine-synthesis-capacity were related to the node importance of the ToM and the eSAD networks in the prediction of cognitive symptomatology.

### Symptom dimensions were differentially predicted by different functional networks

Schizophrenia is a disorder that is commonly proposed with global or widespread brain deficits (10-12). Our network-based predictive modeling, however, revealed specific networks-psychopathology relationships based on the predictive capacity of specific functional systems for patient-specific symptom severity along four dimensions. Two networks, ToM and eSAD, both predicting the cognitive dimension, passed our strict validation steps. The ToM network subserves social cognition while the eSAD network encompasses regions involved in socio-affective processes. These results provide a firm support for the previous findings indicating that the compromised ―social brain‖ development in schizophrenia relates to higher-level cognitive deficits (56,57). Yet it came as somewhat of a surprise that these two networks did not allow a robust prediction of negative or affective symptoms. However, other socio-affective networks, CER, which relates to the reappraisal process of emotional stimuli (58), and MNS, which was proposed to represent the emotional aspect of social cognition (18), were predictive of the positive and affective dimensions, respectively, in the main sample. Hence, it stands to reason that different socio-affective networks would capture individual variance of different symptom dimensions of schizophrenia. Of note, these predictions did not generalize to B-SNIP, potentially due to between-sample differences in clinical characteristics but also highlighting that building models that generalize to completely novel cohorts remains a challenge (59). We also noted that the cognitive dimension was not robustly predicted by any of the cognitive networks employed, though the speech production network showed a trend-level prediction in 10-fold cross-validation (*r*=0.18, *p*=0.061). Previous work showed that task-based functional connectivity yields better predictions of cognition than the networks at rest (60-62). Following this line of thought, cognitive networks engaged during tasks might likewise be more robustly related with the individual variability in cognitive symptoms than intrinsic connectivity patterns.

Overall, we revealed that the connectivity patterns within the ToM network, even at the level of intrinsic brain activity, are robustly predictive of the cognitive status in individual schizophrenia patients. ToM is the cognitive ability of an individual to infer others’ mental states, intentions and believes (63). As these involve complex cognitive processes and considering that the cognitive dimension of schizophrenia psychopathology encompasses symptoms such as ―conceptual disorganization‖, ―lack of insight‖, and ―disturbed abstract thinking‖, it is not unexpected that the ToM network is predictive of the cognitive dimension. Resonating with this finding, ToM deficits, which are prevalent in schizophrenia and a well-established feature and vulnerability marker of this disorder (64), are known to associate with symptoms of disorganization (65). Abnormal neural activation in response to tasks targeting ToM has also been reported in schizophrenia involving temporo-parietal junction as well as prefrontal and temporal regions where the current ToM meta-network is distributed (66,67).

It is interesting to note that the ToM network also predicted negative symptoms in the main sample though this did not generalize to the B-SNIP data. This resonates with the notion that ToM encompasses multiple components (e.g., affective and cognitive)(68) and an impairment in the sub-processes of ToM may manifest as different consequences. And indeed, the subnetworks of ToM predicting negative and cognitive dimensions, respectively, were largely divergent (Figure 3B, Figure S4).

Intriguingly, cognitive symptoms in schizophrenia were also linked to socio-affective processes via the prediction using the eSAD network. Recent studies consistently linked neural activity patterns in the DMN to cognitive abilities (69,70), and the implication of the DMN-derived eSAD network (71) in cognitive processing is hence unsurprising. Our finding moreover resonates well with a literature suggesting that socio-affective factors impact and modulate cognitive performance like working memory and attention in schizophrenia patients (72-74). The use of seemingly non-cognitive psychosocial methods has been proposed as a potential remediation strategy for cognitive deficits in schizophrenia (72), and indeed successfully applied in practice (74). Consequentially, we would hypothesize that the cognitive dysfunctions in schizophrenia might relate to an impaired integration of self- and other-related affective mental processes. Since the interaction of the DMN with other brain regions and networks is more reflective of schizophrenia symptoms than within-DMN connectivity (18,75,76), it is not surprising that the DMN was not predictive, but the eSAD was which comprises regions going beyond the DMN.

### Molecular architecture of the identified networks which robustly predicted cognitive dimension

Cognitive deficits are a lifelong burden for patients with schizophrenia because there are so few effective medications including the mainstay anti-dopaminergic agents (37,77). In line with the notion that dopaminergic dysfunction alone doesn’t account for the whole picture of schizophrenia psychopathology (78,79), here we revealed that the networks and nodes predictive of individual cognitive symptom-load are related to both dopaminergic (32) and serotonergic systems (78). These data extend previous region-of-interest analyses (80,81) to a comprehensive topographical level and are supported by findings that cognitive deficits relate to D_1_ dopamine receptor elevation in schizophrenia (80). The extension to a network-level investigation for the molecular architecturethat is related to symptomatology is important, as schizophrenia patients are characterized by dysconnection between distant brain regions and network connectivity-based analyses can nominate mechanisms of action for the development of novel pharmacological treatments (82). In line with involvement of dopamine functioning in cognitive deficits in schizophrenia (32), the cognitive dimension of psychopathology was associated with the dopamine-synthesis-capacity. Increased dopamine-synthesis-capacity in schizophrenia was not only reported in the striatum (32) but also cortical areas including prefrontal cortex (83,84) where multiple nodes within the ToM and the eSAD networks are located.

Moreover, cognitive symptoms were linked to 5-HTT density at network level. 5-HTT plays a critical role in regulating serotonergic concentration and signaling. Serotonergic dysfunction and 5-HTT polymorphism (85) have been involved in schizophrenia pathophysiology (34,78). Previous postmortem and in vivo PET studies have yielded inconsistent results on the alteration of 5-HTT density in schizophrenia (34), while an over-expression of 5-HTT mRNA in the prefrontal and temporal cortex has been demonstrated (86). We here revealed a potential role of 5-HTT in cognitive deficits via the networks involved in socio-affective processes, resonating with the proposed implication of 5-HTT in the affective domain (87,88). While atypical anti-psychotics including olanzapine and risperidone target also the 5-HT_2a_ serotonin receptor (89,90), their effects on 5-HTT seems to be equivocal (91,92). Interestingly, anti-psychotic drugs, pimavanserin and SEP-363856 have just been introduced that preferentially target serotonin and not dopamine receptors (93,94), suggesting increased focus also on serotonergic pathways.

### Limitations and Considerations

First, the effect sizes for the correlation between the symptom scores and their predictions were moderate. However, despite the clinical complexity of the population and the heterogeneities in scanners and protocols, the effect sizes are similar to the previously reported for predicting, e.g., creativity (95), personality (48), and memory performance (96) in healthy subjects (*r*-values mostly around 0.2-0.35). Since olanzapine-equivalent dosage did not correlate with symptom scores or alter the prediction pattern after additionally controlling for the dosage in cross-validation, medication would be largely a source of random variation in our data and hence make our results more conservative. Second, although glutamatergic dysfunction has been increasingly implicated in schizophrenia neurocognitive deficits (79,97), there are no publicly available in vivo maps reflecting aspects of the glutamatergic system. Finally, within-subject (longitudinal) studies assessing symptoms, rs-FC and receptor densities are needed, though acquisition of molecular imaging data in large samples remains difficult.

Based on rs-FC within different meta-analytic task-activation networks covering a broad range of functional domains and predictive modeling with strict validations, intrinsic connectivity patterns of networks implicated in socio-affective processing was revealed to robustly associate with the cognitive dimension of psychopathology. Our investigation of the molecular architecture of the identified predictive networks implied a potential involvement of 5-HTT serotonin transporter, besides the dopaminergic system, in schizophrenia cognitive symptomatology, possibly providing hints into treatments.

## Acknowledgments

This study was supported by the Deutsche Forschungsgemeinschaft (DFG, EI 816/4-1), the National Institute of Mental Health (R01-MH074457), the Helmholtz Portfolio Theme “Supercomputing and Modeling for the Human Brain”, and the European Union’s Horizon 2020 Research and Innovation Programme under Grant Agreement No. 720270 (HBP SGA1) and 785907 (HBP SGA2). Ji Chen has received a Ph.D fellowship from the Chinese Scholarship Council.

