## Supplemental Materials for "Connectivity patterns of task-specific brain networks allow individual prediction of cognitive symptom dimension of schizophrenia and link to molecular architecture"

### Sample information for each site

#### Main sample

**The Utrecht sample:**

Patients with chronic schizophrenia were diagnosed according to the Diagnostic and Statistical Manual of Mental Disorders, Fourth Edition (DSM-IV) criteria (1) by an independent psychiatrist using the "Comprehensive Assessment of Symptoms and History (CASH)" (2). This study was approved by the Humans Ethics Committee of the University Medical Center Utrecht with written informed consent obtained from the all participants (3).

**The Göttingen sample:**

Patients were recruited from the Department of Psychiatry and Psychotherapy, University Medical Center Göttingen. They met the diagnostic criteria of schizophrenia according to DSM-IV (1) Patients who had substance abuse within the last month, cannabis abuse within the last 2 weeks, past or present substance dependency, somatic or mental disorders that would interfere with the protocol, acute suicidal tendency or an inability to give written consent were excluded (4). Exclusion criteria for subjects in the control group included any DSM-IV diagnosis for the subject or a first-degree relative.

**The Groningen sample:**

Diagnosis of schizophrenia was established based on the DSM-IV criteria (1), confirmed by a Schedules for Clinical Assessment in Neuropsychiatry (SCAN) interview (5). Exclusion criteria included a personal or family history of epileptic seizures, a history of significant head trauma or neurological disorder, the presence of intracerebral or pacemaker implants, inner ear prosthesis or other metal prosthetics/implants, severe behavioral disorders, current substance abuse, and pregnancy (6). The study was approved by the Institutional Review Board of the University Medical Center Groningen.

**The Albuquerque sample (COBRE):**

This dataset was collected and shared by the Mind Research Network and the University of New Mexico funded by a National Institute of Health Center of Biomedical Research Excellence (COBRE; http://fcon_1000.projects.nitrc.org/indi/retro/cobre.html). Patients with schizophrenia were diagnosed based on DSM-IV using the Structured Clinical Interview used for DSM-IV axis I disorders (SCID). Informed consent was obtained from participants at the University of New Mexico. All patients were chronic and with relatively well-treated symptoms by a variety of antipsychotic medications (no medication changes in 1 month). Those patients with a history of neurological disorder, head trauma with loss of consciousness greater than 5 min, mental retardation, active substance dependence or abuse (except for nicotine) within the past year, current use of mood stabilizers, history of dependence on PCP, amphetamines or cocaine, or history of PCP, amphetamine, or cocaine use within the last 12 months were excluded (7). Exclusion criteria included current or past psychiatric disorder, family history of a psychotic disorder in a first-degree relative, history of neurological disorder, head trauma with a loss of consciousness greater than 5 min, mental retardation, recent history of substance abuse or dependence, history of more than one life time depressive episode, history of depression or anti depressant use within the last 6 months, and history of lifetime anti-depressant use of more than 1 year.

**The Aachen-1 sample:**

Patients were diagnosed based on ICD-10 diagnostic criteria (by the corresponding psychiatrist) and further screened with SCID (DSM-criteria). Drug addiction was excluded. Any patients with current drug-use (within the past 6months), neurologic and metabolic disorders were excluded. Symptom severity was assessed using the Positive and Negative Syndrome Scale (PANSS) (8). This study was approved by the ethics committee of the Medical Faculty of the RWTH Aachen University with written informed consent obtained (9).

**The Aachen-2 sample:**

Diagnosis of schizophrenia for the patients in the Aachen#2 sample was established using the ICD-10 diagnostic criteria (code F20.X). Symptom severity in schizophrenia patients was measured using the PANSS (8). Exclusion criteria for all subjects were 1) current substance or alcohol abuse or any diagnosed substance abuse or addiction in the past (ICD-10: F10.1 or .2 - 19.1 or .2), 2) severe medical conditions such as chronic or acute diseases (i.e., infections, allergies), 3) general contraindications to MRI scanning, 4) gross morphological changes on MRI such as cerebral atrophy, hydrocephalus or previous injury, and 5) incomplete scanning. Subjects treated with benzodiazepines were not included, except the drug was at least in its second elimination half-life period after intake. The ethics committee of the Medical Faculty of the RWTH Aachen University approved this study with written informed consent obtained.

**The Lille sample:**

Patients were diagnosed with schizophrenia according to the DSM-IV-TR criteria (10). All patients routinely presented frequent (more than 10 per day) and resistant hallucinations as evaluated with item P3 of the PANSS. The exclusion criteria included the presence of an Axis-II diagnosis, secondary Axis-I diagnosis, neurological or sensory disorder, and a history of drug abuse, which was based on a clinical interview and urine tests that were administered at admission. The study was approved by the local ethics committee (CPP Nord-Ouest IV, France). Written informed consent from each patient was obtained (11).

#### The B-SNIP sample for validation

Subjects with schizophrenia in the independent, validation sample were recruited as part of the Bipolar-Schizophrenia Network on Intermediate Phenotypes (B-SNIP) Consortium study which used identical diagnostic and clinical assessment techniques with similar recruitment approaches at multiple sites (Baltimore, Chicago, Dallas, Detroit, and Hartford). Detailed information on the entire study sample is provided elsewhere (12). These clinical patients were diagnosed using the Structured Clinical Interview for DSM-IV Axis I Disorders, Patient Edition (SCID-I/P) and were stably medicated outpatients. The study protocol was approved by the institutional review board at each local site and written informed consent was obtained from each of the volunteers. For our current study purpose, only schizophrenia patients with complete PANSS and fMRI data were included. Among the four sites (Baltimore, Dallas, Detroit, and Hartford) data we retrieved, the Dallas site was scanned using a Philips machine such that the details on the order of each slice scanned within a EPI volume were not stored in the *Header* of the resting-state images and was hence excluded, because inaccurate slice timing correction will lead to a biased estimation of rs-FC. The Detroit site was also not included due to the small sample size (<10) retained after sample curation and quality control. Resultantly, the dataset (in total 117 schizophrenia patients) pooled from the Hartford site (40 patients) and the Baltimore site (77 patients) was used as an independent validation of our predictive modeling. The sample size was considered to be sufficient. That is, if the highest effect size of 0.31 (i.e., the correlation r: observed dimensional symptom scores vs. their out-of-sample predictions) which was identified in 10-fold cross-validation within the main sample could be replicated in the validation sample, the minimal sample size for detecting a significant correlation of 0.31 at an α-level of 0.05 (two-tailed) and a power of 80% would be 76 subjects. Our validation sample includes 117 subjects, which still has a power of 70% for detecting a statistical significance at the α-level of 0.05 (one-tailed) even at a lower effect size of 0.2. Therefore, the sample size of 117 was deemed sufficient for validation. Power analysis was performed using the G*Power software (https://www.psychologie.hhu.de/arbeitsgruppen/allgemeine-psychologie-und-arbeitspsychologie/gpower.html) (13). Demographic and image scanning details for both the main and the B-SNIP samples are provided in Tables S1, S2, S4 and S5.

### Supplementary Materials and Methods

#### MRI data acquisition and preprocessing

Specific scanning parameters for high-resolution T1-weighted anatomical and resting-state fMRI images were provided in Tables S4 and S5. Image preprocessing was done in Statistical Parametric Mapping software (SPM12; https://www.fil.ion.ucl.ac.uk/spm) and Computational Anatomy Toolbox (CAT12; https://www.neuro.uni-jena.de/cat) and is detailed elsewhere (14). In brief, for resting-state modality, the first four volumes from all fMRI scans were discarded. Then, the DVARS metric (15) was employed by calculating the voxel-wise BOLD signal intensity change between one frame (timepoint) and it's backward to detect and remove the patients with excessive movements. The DVARS metric was scaled by dividing by the median brain intensity and then multiplying by 1000 to approximate the magnitude reported in Power *et al.* (15), i.e., 10 units of DVARS refer to 1% BOLD signal change. As in our previous study which used the same patient cohort for imaging analyses (14), any patient with a DVARS larger than 50 (i.e., 5% BOLD signal change) was removed from subsequent predictive modeling. The cutoff of DVARS=50 is roughly equivalent to a framewise displacement of 0.5mm commonly used in the literature (16). Afterward, all of the images were slice timing corrected (17) and head motion corrected in SPM12, and the derived six motion parameters were used as motion regressors. The motion-corrected EPI data were normalized to MNI152 space by using an EPI template in SPM12 with a 4 x 5 x 4 basis set to alleviate overfitting (18). The normalized EPI images were resampled to an isotropic voxel size of 2mm. The high-resolution T1-weighted structural images were preprocessed in CAT12. The resultant partial volume image for each patient containing the three grey matter, white matter (WM) and CSF tissue types was used as masks for extracting global mean white matter (WM) and CSF signals. Here a quality control analysis was conducted by using the "check sample homogeneity" module in CAT12 to filter out those subjects with poor segmentation quality in structural images as this might lead to an inaccurate estimation of the WM and CSF signals. Twenty-four head motion parameters (the six head motion parameters of roll, pitch, yaw, translation in three dimensions, their first temporal derivatives, and quadratic term signals), together with the non-neuronal components of the extracted total WM and CSF signals were regressed out from the overall BOLD signals (19). As still a controversial step in resting-state fMRI preprocessing (20,21), we did not regress out global mean signals in our analysis. Global signal regression was moreover reported to obscure an effect predictive of symptoms in schizophrenia (22). Finally, band-pass filtering was performed on the data to restrict frequencies between 0.01 and 0.08 Hz.

After estimating subject-wise head movement and assessing T1-image segmentation, 16 patients in the main sample were found to have excessive head motions and five patients were with poor tissue segmentation quality and hence these 21 patients were removed from the subsequent analyses. In the validation B-SNIP sample, one patient with schizophrenia was excluded from the Hartford site due to an excessive head-motion, while 16 patients in the Baltimore site were excluded (15 patients had excessive head-motions and one patient with bad tissue segmentation in the resulting *T1* partial volume image). After filtering out the in total 17 schizophrenia patients in B-SNIP, 39 patients in the Hartford site and 61 patients in the Baltimore were retained for validation analysis. In the remaining patients, age did not correlate with any of the four symptom dimensions in the main (all $p$-values>0.25; Pearson correlation analysis) and the B-SNIP samples (all $p$-values>0.20). No gender differences were observed for the scores of the four symptom dimensions within the main sample (all $p$-values>0.09). While in the B-SNIP sample, male patients (2.43±1.67) showed significantly lower ($p$=0.018) affective dimensional-scores than female patients (3.31±1.69). Scores for the other three symptom dimensions did not show any gender differences within the B-SNIP sample. Age and gender were both adjusted in our predictive modeling to avoid (any) possible contributions from them.

#### Predicting individual symptom dimensional-scores from within-network rsFC patterns

##### Detailed methodology for relevance vector machine

To approach the prediction problem, we employed a relevance vector machine (RVM) (23) as implemented in the SparseBayes package (http://www.miketipping.com/index.htm) to achieve multivariable regression so that continuous target variables can be predicted based on set of features (i.e., exploratory variables). The RVM refers to a specialization of the general Bayesian framework which has an identical functional form to the support vector machine (SVM) 24). In SVM, a separating hyperplane is computed based on the feature space learned from the training data by maximizing the margin between the two groups. The SV regression (SVR) extends the binary outputs from SVM to achieve an estimation and prediction of continuous variables. Similar to the margin generated in SVM classification, the regression line of SVR is surrounded by a tube. Unlike SVM/SVR, RVM is embedded in a probabilistic Bayesian framework which substitutes the margin term in SVM by a prior distribution over the parameters. Specifically, an explicit zero-mean Gaussian prior is imposed to avoid severe overfitting associated with the maximum likelihood estimation of the model weights (25,26). Sparsity can be achieved in RVM through sparse modeling with no additional penalty terms needed to shrink the predictor coefficients as the posterior distributions of many of the estimated weights are already towards zero. By computing the predictive distribution, the target value of a previously unseen input vector can be predicted from the trained model. RVM is free from constraints on the kernel functions (such as the Mercer's condition that required by SVM) and utilizes dramatically fewer basis functions. Moreover, the control parameters in RVM can be automatically estimated by the learning procedure itself (i.e., no hyperparameters that need to be tuned), which thus, exerts enhanced efficiency comparing to a classical SVM/SVR.

#### Leave-one-site-out cross-validation

Leave-one-site-out cross-validation, i.e., training on the data without one site and then testing on the left-out site to assess the generalizability of our predictive model to new sites with different scanners, was performed on the main sample. Specifically, in each leave-one-site-out realization, we left each single site out, trained models on the other sites, predicted the left-out site. The process was repeated until each site had been left out once, and, finally, we calculated the Pearson's correlation coefficient between the actual (confound-adjusted) symptom dimensional-scores and their predictions. As in previous studies (27,28) and in keeping with our 10-fold process, confounding effects of age, gender and head motion on both the symptom dimensional scores and within-network rsFC in the left-out test site and the remaining six sites (training set) were adjusted using the regression weights estimated from the training sites.

For the validation analysis in B-SNIP, we likewise adjusted the effects of age, gender, and head motion on both the symptom dimensional-scores and rs-FC features using the regression weights estimated from the main sample (i.e., the training set) (28). Site effects were adjusted by fitting a linear model within these two sites B-SNIP data (29).

#### Permutation tests for assessing the significance of cross-validated-correlations

The folds in 10-fold cross-validation are not completely independent, the number of degrees of freedom (DOFs) is thus overestimated and using parametric statistical tests here to derive the p values is problematic (30,31). Therefore, we implemented iterative permutation tests on the actual data to estimate the significance of the cross-validation-based correlations. In each realization of the permutation test to establish the null distribution for network-based prediction: 1). the symptom dimensional scores are shuffled randomly between subjects while keeping everything else exactly the same, 2). the same stratified 10-fold cross-validation was applied to predict the shuffled scores and the correlation coefficient between the shuffled ratings and their out-of-sample predictions was recorded. Afterward, we repeated the above two steps for 1000 times. The whole process allowed us to create an empirical distribution of the chance (i.e., based on the shuffled dimensional scores) correlation to compare the true (i.e., based on the original ratings) correlation coefficient against. If the correlation coefficient of non-permutated scores (original ratings) exceeds those obtained from the all 1000 permutation tests, indicating a statistical significance of $p = 0.001$ (i.e., right-tailed). The same iterative permutation tests were implemented to derive the significance for the leave-one-site-out cross-validated correlation since the sites used for model training are not completely independent of the test sites in the seven leave-one-site-out experiments).

For the validation analysis in the independent B-SNIP sample, *p* values were derived by using standard parametric statistical tests because the samples used for model training and the sample employed for model testing were completely independent and the assessment of statistical significance is hence free from the DOF-overestimation issue.

#### Details for the density maps of receptors/transporters from prior molecular imaging studies

As detailed in our prior study (32), the density estimates of gamma-aminobutyric acid (GABAa) and dopamine transporter (DAT) were obtained from flumazenil positron emission tomography (PET) and single photon emission tomography (SPECT), respectively. Here in brief:

***DAT***

Baseline DAT-SPECT data of 174 healthy elderly volunteers (mean age±SD: 61±11 years, 109 males) were extracted from the Parkinson's Progression Marker Initiative database (PPMI, www.ppmi-info.org/) (33). Written informed consent was obtained from all subjects. The study was approved by Institutional Review Boards/Independent Ethics Committees. A mean image was computed from the preprocessed data in MNI space with a Gaussian kernel of 8mm FWHM. DAT density estimates were extracted from this mean image. Of note, previous studies showed that in vivo DAT density in the brain as assessed by SPECT declines with age (mostly linear) which is typically up to 10% per decade year for some regions (e.g., caudate and putamen) (34-36). However, that should be less of a concern with respect to our current network-based analysis relying on the relative ratio of regions to each other which is robust to age. This is because the expression of DAT in some specific brain regions e.g., the basal ganglia (and to some little extent the prefrontal cortex) is in very high amounts and hence the relative differences between these DAT-rich areas and other DAT-poverty brain regions (e.g., thalamus) (36) are still fairly large. That is, these observation and relative order of brain regions with respect to each other remains stable throughout a healthy life span. Also, here we used Spearman's rank correlation which assesses monotonic relationships. .

***GABAa***

Dynamic [11C]flumazenil PET scans were acquired from 6 healthy volunteers with full arterial blood sampling for quantitative compartmental modeling (37). PET images were reconstructed into 20 frames using filtered back projection. Voxel level spectral analysis (38) with 100 logarithmically distributed orthogonal basis functions between 0.0008 and 1 s-1 was performed to create parametric maps of total distribution volume (VT), with 2.09x2.09x2.42 mm resolution. These individual volumes of distribution maps calculated as the summed integral of the peaks after spectral analysis were used as individual GABAa density estimates and were then normalized to MNI space. This study was approved by a NHS Research Ethics Committee, the Administration of Radioactive Substances Advisory Committee and local NHS Research and Development.

***D2/3 (https://datadryad.org/resource/doi:10.5061/dryad.rc073)***

The radiotracer [11C]raclopride binding in striatal subregions, the thalamus and the cortex was investigated using the bolus-plus-infusion method and a high resolution PET (39). Seven healthy male volunteers underwent two PET [11C]raclopride assessments, with a 5-week retest interval. Spatial resolution in the reconstructed PET images varies in radial and tangential directions from ~2.5 to 3 mm and in axial directions from 2.5 to 3.5 mm in the 10-cm field of view covering the brain. $D_{2/3}$ receptor density was quantified as binding potential using the simplified reference tissue mode tissue compartmental modeling (SRTM) (40,41). The cerebellum which is devoid of $D_{2/3}$ receptors (42) was chosen as the reference tissue. For voxel-level model fitting, they used a linearized model using a basis-function approach (43) implemented also in in-house software (http://www.turkupetcentre.net/programs/doc/imgbfbp.html). The absolute variability and intraclass correlation coefficient values demonstrated good test-retest reliability. We accessed to the baseline session of scanning and got the group average whole-brain map with voxel-wise D2/3 density estimates. The study protocol was approved by the Ethics Committee of the Hospital District of Southwestern Finland. Written consent was obtained from each volunteer.

***D1***

Thirteen healthy volunteers (7 female, age 33 ± 13 yrs) underwent 90-min emission scans, each after 90-s bolus injection of 486 ± 16 MBq [11C]SCH23390, on two separate days within 2-4 weeks using a PET/MRI system (44). This study was approved by the local ethics committee (registration number 083/11) and the German Federal Office for Radiation Protection (number Z5-22461/2-2012-003). Informed consent was obtained from all participants. Motion correction was performed with Statistical Parametric Mapping (SPM8, Wellcome Department of Cognitive Neurology, London, UK). Individual MRI T1-weighted MPRAGE data sets and the related, already-fused PET data of each subject were spatially reoriented onto a standard brain data set similar to the Talairach space, reconstructing the images to 128×128×64 voxels with dimensions of 1.7×1.7×2.5 $mm^3$. Parametric images of binding potential (BPND) were generated in PMOD (version 3.5, PMOD Technologies, Zurich, Switzerland) from the PET data by the multi-linear reference tissue model with two parameters (MRTM2) and the cerebellar cortex as the receptor-free reference tissue (45). The BPND maps were used in our present spatial correlation analysis as a reflection of D1 receptor availability.

***$^{18}$F-DOPA (https://www.nitrc.org/projects/spmtemplates/):***

Participants' dopamine synthesis capacity was measured by using [18F]DOPA PET. The map we used for local DSC estimates was the 18F-DOPA template in SPM. Data acquisition and template construction were detailed in the original publication (46) and briefly as follows: Brain PET images were acquired with 18F-DOPA to 12 control subjects (6 males and 6 women aged 55.1 ± 16.6 years) without evidence of nigrostriatal degeneration. PET experiments were performed on a CT scanner Siemens Biograph 16 PET/CT, which provides a 2.0 mm FWHM of the FOV. One hour before the injection of the established dose of 18F-DOPA, 150 mg of carbidopa was administered by oral to block the enzymatic activity of the DOPA decarboxylase. The PET images acquisition started 90 minutes after intravenous injection of the radiotracer using 222 MBq. The emission PET data were acquired for 20 minutes in 3D mode, after a brain CT scan in spiral mode at 120 kVp and 160 mA with the CARE program Dose 4D. The raw data were reconstructed using the OSEM algorithm with 4 iterations, 8 subsets, all-pass filter and a matrix resolution of 128 × 128. All images were transformed from DICOM format to NIfTI using the dcm2nii function in MRIcron (http://www.mccauslandcenter.sc.edu/mricro/mricron/). Then, the baseline PET scans were spatially normalized to a common anatomical space using a T1-weighted structural MRI template as reference in software SPM8 (SPM; Welcome Department of Cognitive Neurology. London, UK). To avoid the intrinsic asymmetries presented in the recruited sample, the left-right hemisphere flipped (i.e., mirrored) images of the original PET scans for each of the 12 patients were obtained, resulting in total 24 brain PET images for subsequent template construction. Afterward, an intensity normalization procedure was performed on the PET images, which resulted in each voxel has a value between 0 and 1. For a precise alignment to the standardized anatomical space of MNI152 (MNI; http://www.bic.mni.mcgill.ca), the resultant maps were resampled with a bounding box of 90 × 109 × 91 and an isotropic voxel size of 2mm. Finally, the 24 PET images were averaged to form the template, and the value for each voxel in the template was the group mean intensity-normalized value with a Gaussian filter step applied. The authors declared that the procedures conformed to the ethical standards of the responsible human experimentation committee.

***Serotonergic receptors and serotonin reuptake transporter***

For serotonergic system, including the three serotonin receptors of 5HT1a, 5HT1b, 5HT2a and serotonin reuptake transporter 5-HTT, the density estimates were derived from a multi-center PET study with different radiotracers (47). A total of 95 healthy subjects (mean age= 28.0±6.9 years, range= 18-54, 59% males) were included in this multicenter PET to map the serotonergic receptors and transporter *in-vivo*. All subjects were physically healthy and life-time naïve for psychotropic drugs. All participants gave written informed consent according to the procedures approved by the local Ethics Committees at the Medical University of Vienna, the Medical Faculty of the University of Düsseldorf and the Yale School of Medicine Human Investigation Committee.

For assessing ***5HT1a,*** the [carbonyl-$^{11}$C]WAY-100635 was used as the radioligand. PET scans were conducted with a GE Advance PET scanner (General Electric Medical Systems, Milwaukee, Wisconsin) with a spatial resolution of 4.36 mm full-width at half maximum (FWHM) at the center of the FOV (35 slices). Details in image acquisition and reconstruction are described elsewhere (48).

For ***5HT1b,*** [$^{11}$C]P943 PET scans were acquired for 120 min on an HRRT PET scanner (207 slices, resolution less than 3 mm full-width at half maximum in 3D acquisition mode). PET image acquisition and image reconstruction were performed as described previously (49).

For the ***5-HT2a*** receptor, a highly selective radioligand of [$^{18}$F]altanserin was used. PET measurements were performed in 3D mode on a Siemens ECAT EXACT HR+ scanner (Siemens-CTI, Knoxville, TN, USA; 63 slices; full-width of half maximum 5.8, 5.8, 6.6 mm (x, y, z) at 10 cm from the central axis). Tracer application according to a 2-min bolus plus constant infusion schedule (KBol= 2.1 h), venous blood sampling, metabolite correction of the plasma input function, PET image acquisition and reconstruction were conducted according to the previous publication (50).

For the *in-vivo* quantification of ***5-HTT*** serotonin transporter, [11C]DASB is used as the radioligand which is high affinity and selectivity to 5-HTT (Wilson et al., 2000, 2002). PET scans were obtained from a GE Advance PET scanner (General Electric Medical Systems, Milwaukee, Wisconsin) with a spatial resolution of 4.36 mm full-width at half maximum (FWHM) at the center of the FOV (35 slices).

Apart from the preprocessing of 5-HT2a scans which was performed at the Research Centre Jülich using SPM2 (50), the raw PET scans for other receptors and the 5-HTT transporter were preprocessed using SPM8 (http://www.fil.ion.ucl.ac.uk/spm/software/spm8/) and the motion correction was carried out by co-registration of each frame to the mean of the subjects' motion-free frames. Dynamic PET scans were normalized onto tracer-specific templates in MNI stereotactic space by computing the transformation matrices of individual PETADD (sum over all time frames) and subsequent application to dynamic scans. Ligand-specific templates were created following the approach introduced previously (51) and provided mean values for each voxel. The original PET images were further spatially smoothed with an isotropic 8 mm Gaussian kernel. Finally, the quality of spatially normalized images was visually inspected where necessary.

The binding potential (41) was calculated using the kinetic modeling tool PKIN as implemented in PMOD (PMOD Technologies Ltd, Zürich, Switzerland) to denote receptor density. Cerebellum was used as the reference region. For quantification of [carbonyl-11C] WAY-100635, [11C]P943 and [11C]DASB binding, the author used the "multilinear reference tissue model" (MRTM/MRTM2) as described previously (45) to calculate the $BP_{ND}$, while the [18F]altanserin scans were parameterized on the basis of the cerebellum ($C_{Reference}$) and the plasma activity concentration attributable to parent compound ($C_{Plasma}$) using the following equation: $BP_P=(C_{ROI}-C_{Reference})/C_{Plasma}$ with radioactivity concentrations averaged from 120 to 180 min p.i. (52). [18F]altanserin binding potentials were read out from parameterized maps.

Overall, for comparability, the maps of density estimates obtained from aforementioned multi-tracer molecular imaging studies, in MNI152 space, were linearly rescaled to a minimum of 0 and a maximum of 100 and were resampled to an isotropic 2mm spatial resolution (original resolutions: 1.7-6.6mm) as in our fMRI data. The closest between-node distance within the two robustly predictive networks, theory-of-mind and extended socio-affective default, is 12mm, and thus the nodal density estimates for the investigated receptors/transporters were differentiable in the molecular data.

#### Assessing statistical significance for spatial correlation analysis

We implemented a spatial permutation testing to assess the statistical significance for the spatial correlation between network nodes and receptor/transporter densities calculated for these nodes. That is, we generated 1000 random networks by re-distributing the nodes throughout the grey matter with the same number of nodes as in real network while preserving the between-node distance (±6mm tolerance). Nodal receptor/transporter densities were extracted from these simulated (random) networks which were then correlated with the node importance scores for the real network. This allowed us to construct a null with 1000 (chance-level) correlations based on a set of randomized topographic configuration of networks. Finally, the true correlation based on the nodal receptor/transporter densities extracted from the real network was compared with the null distribution to derive the significance (lowest $p$=0.001). If the true correlation obtained from the real network exceeds the 95% percentile of the null, indicating a statistical significance for the true correlation against a (pseudo)-random placement of nodes within the grey matter.

Several metrics were additionally employed to assess the property of the generated random networks, including:

1) between-node distances within each random network;

2) distance between random networks;

3) distance between the original (real) and the random networks.

Metrics 2) and 3) allowed to check if the random networks are adequately different from each other and from the real network.

As shown in Figure S3, these normally distributed histograms demonstrated that our simulated random networks well reflected possible spatial configurations within the grey matter of the entire brain, as these random networks were sufficiently different from, but neither too far or too close to, each other and the real networks.

**Table S1.** **Clinical characteristics of the main, international schizophrenia sample for each site**

| | Aachen-1 | Aachen-2 | Albuquerque (COBRE) | Göttingen | Groningen | Utrecht | Lille | Total | *P*-value[1] |
|---|---|---|---|---|---|---|---|---|---|
| ***N*** | 13 | 10 | 47 | 32 | 22 | 10 | 13 | 147 | |
| Illness duration | 7.93± 8.52 | 13.11 ±10.93 | 16.66 ±12.21 | 7.03 ±7.59 | 7.64 ±7.23 | 8.5 ±7.23 | 12.38±6.06 | 11.23 ± 10.31 | **<0.001** |
| Gender (Male/Female) | 10/3 | 5/5 | 36/11 | 26/6 | 13/9 | 5/5 | 8/5 | 103/44 | 0.178 |
| Age | 35.07±11.15 | 34±9.77 | 37.72±13.9 | 32.28±9.94 | 34.05±12.83 | 33.3±8.69 | 32.77±8.45 | 34.77±14.71 | 0.537 |
| ***Antipsychotic treatment*** | | | | | | | | | |
| Typical antipsychotics | 0 | 0 | 4 | 0 | 1 | 1 | 1 | 7 | |
| Atypical antipsychotics | 13 | 10 | 41 | 26 | 19 | 4 | 10 | 123 | |
| Both atypical and typical antipsychotics | 0 | 0 | 2 | 5 | 0 | 0 | 1 | 8 | |
| Missing/None | 0 | 0 | 0 | 1 | 2 | 5 | 1 | 9 | |
| olanzapine -equivalent[2] | 21.72±10.05 | 18.92±12.87 | 14.84 ±10.96 | 25.06 ± 11.49 | 14.55 ± 8.31 | 17.10 ± 12.42 | 26.24±21.21 | 18.81 ± 11.60 | **0.001** |
| ***Scores on the Four Dimensions of PANSS*** | | | | | | | | | |
| Negative | 1.78±1.68 | 5.99±3.94 | 2.41±2.05 | 1.93±1.59 | 2.30±2.20 | 3.69±2.36 | 5.59±2.96 | 2.92±2.53 | **<0.001** |
| Positive | 2.99±2.23 | 4.47±2.97 | 2.99±2.03 | 1.61±1.54 | 4.27±2.32 | 4.91±1.44 | 6.45±2.12 | 3.41±2.44 | **<0.001** |
| Affective | 2.65±1.79 | 6.70±3.29 | 3.12±2.19 | 2.76±1.60 | 3.01±2.11 | 3.33±1.73 | 5.12±3.10 | 3.40±2.38 | **<0.001** |

| | | | | | | | | | |
|---|---|---|---|---|---|---|---|---|---|
| Cognitive | 1.35±1.26 | 5.91±2.61 | 2.12±1.22 | 2.10±1.35 | 2.29±1.84 | 2.69±1.43 | 5.90±2.30 | 2.67±2.08 | **<0.001** |
| ***PANSS subscales*** | | | | | | | | | |
| Positive | 15.36±6.70 | 17.11 ± 5.75 | 14.53 ± 5.07 | 11.72 ± 3.52 | 16.59 ± 5.18 | 17.2 ± 2.78 | 22.08±4.70 | 15.31 ± 5.52 | **<0.001** |
| Negative | 11.14±3.90 | 24.00±8.00 | 14.08±4.56 | 12.75±4.22 | 14.45±4.97 | 17.50±5.72 | 22.15±6.07 | 15.02±6.05 | **<0.001** |
| General | 25.29±6.71 | 49.44±13.09 | 28.34±8.48 | 27.56±5.88 | 29.55±8.56 | 30.50±8.95 | 44.15±14.35 | 30.90±10.98 | **<0.001** |
| Symptom severity (Total score) | 51.79±15.44 | 90.56±22.89 | 56.98±13.79 | 52.03±10.36 | 60.59±16.24 | 65.20±14.75 | 88.38±23.86 | 61.33±19.58 | **<0.001** |

*Note:* Data are mean ± SD. N: number of subjects per research site; PANSS, Positive and Negative Symptom Scale; [1]Statistical comparison between sites was conducted using either one-way analysis of variance (ANOVA) or chi-square test where appropriate. [2]Dosage in mg/day.

**Table S2. Demographic and clinical characteristics of schizophrenia patients retrieved from the B-SNIP database**

| **Characteristics** | Hartford (N=40) | Baltimore (N=77) | *p*-value |
|---|---|---|---|
| ***Demographic*** | | | |
| Age (years)[a] | 30.4 (10.98) | 37.25 (12.641) | **0.004** |
| Gender (male/female) | 29/11 | 56/21 | 0.979 |
| Illness during (years)[b] | 7.62 (7.98) | 15.19 (11.77) | **0.001** |
| ***PANSS*** | | | |
| Positive[c] | 15.03 (5.13) | 14.69 (6.19) | 0.768 |
| Negative | 14.78 (6.69) | 15.40 (5.24) | 0.578 |
| General[d] | 30.68 (8.39) | 25.94 (6.12) | 0.001 |
| Illness severity (Total PANSS)[e] | 60.48 (17.91) | 56.03 (13.74) | 0.138 |
| ***Loadings on the dimensions of PANSS*** | | | |
| Negative | 2.74 (2.58) | 2.85 (2.05) | 0.798 |
| Positive | 3.27 (2.47) | 3.19 (2.58) | 0.882 |
| Affective | 3.25 (1.70) | 2.19 (1.61) | **0.001** |
| Cognitive | 2.89 (1.93) | 2.55 (1.65) | 0.310 |

*Note:* Data are mean (SD). *p*-values in bold indicate a significance of $p < 0.05$. Except for gender, which was based on chi-square test, other statistics were based on two sample t-test.

**Table S3. Meta-analytic functional brain networks, domains and their implications in schizophrenia**

| Description of meta-networks | | | | | Involvements in schizophrenia | |
|---|---|---|---|---|---|---|
| ***Domain*** Network (Abbr.) | **Linked processes** | **Experiments/tasks/contrasts for deriving the networks** | **Number of nodes** | **Source publication** | **General summary** | **References** |
| ***Affective*** | | | | | | |
| Emotional scene and face processing(EmoSF) | Perception of emotional scenes and faces | Discrimination of emotional faces and scenes from neutral | 24 | Sabatinelli, (53) | Identifying and discriminating among different facial expressions are impaired in schizophrenia patients. Worse performance and hyper/hypo-activation in frontal and limbic (e.g., amygdala and hippocampus) regions in response to facial emotion recognition, as well as altered functional subnetwork during emotional face processing were reported. | Cao *et al.* (54)<br>Edwards *et al.* (55)<br>Gur *et al.* (56)<br>Phillips *et al.* (57)<br>Adolphs *et al.* (58) |
| Reward-related decision making (Rew) | Reward value-based preferences for possible options, selecting and executing actions, and evaluating the outcome | Convergence across reward valence and decision stages | 23 | Liu *et al.* (59) | Decision making is disrupted in schizophrenia that the patients are inability to properly estimate reward value, which has been related to severity of negative and cognitive symptoms, and linked to prefrontal GABAergic dysfunction. Increased activations were observed in anterior insula, putamen, and frontal sub-regions in response to reward outcomes. | Piantadosi *et al.* (60)<br>Tikàsz *et al.* (61)<br>Kim *et al.* (62)<br>Collins *et al.* (63)<br>Gold *et al.* (64,65) |
| Cognitive emotion regulation (CER) | Reappraisal of emotional stimulus | Reappraise > naturalistic emotional responses | 14 | Buhle *et al.* (66) | Individuals with schizophrenia display emotional regulation abnormalities and cognitive control deficits which tend to increase negative emotion and cause prepotent response of unpleasant scenes via reappraisal. | Strauss *et al.* (67);<br>Sullivan *et al.* (68) |
| ***Social*** | | | | | | |
| Empathy | conscious and isomorphic experience of somebody else's affective state | "feel into" affect-laden social situations > watched or listened passively | 22 | Bzdok (69) | Empathy deficits are presented in schizophrenia which would lead to social dysfunction. Fronto-temporal functional connectivity was related to cognitive empathy and experiential negative symptoms. Reduced cortical | Bonfils *et al.* (70)<br>Singh *et al.* (71)<br>Abram *et al.* (72)<br>Massey *et al.* (73) |

| | | | | | | |
|---|---|---|---|---|---|---|
| | | | | | thickness in empathy-related neural regions (e.g., mPFC, aMCC, and insula) was demonstrated. Reduced activation in fusiform gyrus, lingual gyrus, middle and inferior occipital gyrus was found in schizophrenia patients during empathy task and reduced grey and white matter volumes were observed in these same brain areas. | |
| Mirror neuron system (MNS) | mental imitation (i.e., ‘mirroring’) of others' nonverbal expression (e.g., actions and behavior) | Action observation ∩ action imitation | 11 | Caspers *et al.* (74) | Dysfunctional mirror neuron activity (MNA) has been associated with diverse symptoms (negative, affective) in schizophrenia and abnormal (including both increased and decreased) MNA have been found in the patients | Mehta *et al.* (75,76); Horan *et al.* (77); Pridmore *et al.* (78) |
| Theory-of-mind (ToM) | the cognitive ability of an individual to 'infer the mental states of others' | ToM > non-social baseline | 15 | Bzdok *et al.* (69) | ToM is impaired in schizophrenia, serving as a well-established feature and vulnerability marker of this disorder. The neurobiological basis of ToM deficits has also been indicated previously including findings of abnormal brain activations (temporoparietal junction, middle prefrontal/inferior frontal cortex, posterior cingulate cortex [PCC] and temporal area) in response to tasks targeting ToM and altered brain functional connectivity has also been observed in some DMN regions in schizophrenia. Multiple schizophrenia symptoms (e.g., positive, negative and disorganized) have been associated with ToM deficits. | Bora and Pantelis, (79); Fretland *et al.* (80); Benedetti (81); Shamay-Tsoory (82); Mothersill *et al.* (83); Das *et al.* (84) |
| ***Task-deactivation and interacting*** | | | | | | |
| Extend socio-affective default (eSAD) | A general default mode of socio-affective processing | Regions within the DMN that are consistently found to relate with socio-affective processing[a], together with their intimately coupled regions identified by MACM and ALE | 12 | Amft *et al.* (85) | Impaired social functioning is associated with cognitive (e.g., working memory) deficits in schizophrenia. Abnormal activation in PCC in response to socio-affective related tasks was co-varied with PCC-vmPFC functional connectivity at rest and correlated with negative symptoms. Abnormal social-affective processing has also | Ebisch *et al.* (86) Hendler *et al.* (87) Park *et al.* (88) |

| | | | | | | |
|---|---|---|---|---|---|---|
| | | | | | been related to disturbed functional cohesion within social-affective affiliation networks. | |
| Default mode network (DMN) | Active at rest or during passive rest and mind-wandering that relates to a variety of functions including self-reference, autobiographical information, theory-of-mind and episodic memory. | Contrasts that were coded as a Deactivation (e.g., Control - Task) using a Low-Level Control (strictly defined as either resting or fixation conditions) across a wide range of paradigms (i.e., task-independent deactivations) | 9 | Laird *et al.* (89) | The DMN has been frequently investigated and consistently reported as abnormal in schizophrenia, both structurally and functionally (e.g., reduced grey matter volume, increased and decreased deactivations and functional hyperconnectivity) in e.g., medial prefrontal cortex, anterior/posterior cingulate cortex, and middle temporal gyrus were revealed and have been associated with negative symptoms and cognitive deficits. | Garrity *et al.* (90)<br>Hu *et al.* (91)<br>Jia *et al.* (92)<br>Pomarol-Clotet *et al.* (93)<br>Du *et al.* (94) |
| ***Executive*** | | | | | | |
| Vigilant attention (VigAtt) | Maintaining stable and focused attention | Tasks posing only minimal cognitive demands on the selectivity and executive aspects of attention for more than 10s | 16 | Langner, (95) | Deficits of attention are common in schizophrenia, and abnormal functional brain response to attentional tasks was reported in the frontal cortex, postcentral gyrus, medial temporal lobe and cerebellum. Also, sustained attention was found to correlate with negative symptom severity. | Eyler *et al.* (96)<br>O'Gráda *et al.* (97) |
| Cognitive action control (CogAC) | Supervisory control for the suppression of a routine action in favor of another, non-routine one | ALE coordinate-based meta-analysis on stroop-task, spatial interference task, stop-signal task and go/no-go tasks | 19 | Cieslik *et al.* (98) | Impaired action control was frequently reported in schizophrenia which may influence performance in a wide variety of cognitive domains and are associated with deficits in prefrontal-based control network particularly in (dorsolateral prefrontal cortex as well as premotor, ACC and thalamus. | Reuter *et al.* (99)<br>Braver *et al.* (100)<br>Barch (101)<br>Minzenberg *et al.* (102) |
| Extend multi-demand network (eMDN) | Performance of executive functioning across multiple demands | Using regions of the MDN[b] as seeds for whole-brain resting-state and MACM analyses. The eMDN was then delineated by identifying | 17 | Camilleri *et al.* (103) | The general executive cognition comprises multiple processes that are related but not limited to action/inhibitory control, attention, working memory, and reasoning, which all have been implicated as abnormal in schizophrenia and altered neural activations were | Giraldo-Chica *et al.* (104)<br>Rubia *et al.* (105)<br>Langdon *et al.* (106)<br>Ramsey *et al.* (107) |

| | | | | | | |
|---|---|---|---|---|---|---|
| | | regions in which the consensus connectivity maps of at least half of the seeds overlapped | | | consistently found in anterior cingulate cortex, dorsolateral prefrontal and thalamus in the response to executive-related tasks. | Minzenberg *et al.* (108) |
| Working memory (WM) | A limited resource that is distributed flexibly among all items to be maintained in memory | Consistently activated during all WM contrasts/experiments (mainly n-back, Stenberg, DMTS, delayed simple matching) | 22 | Rottschy, (109) | As a cardinal cognitive symptom that may underlie many other cognitive deficits and symptoms, impaired WM is a persistent, disabling feature of schizophrenia, which has been frequently associated with (dorso/ventro lateral) prefrontal dysfunction (e.g., abnormal neural activation and dopamine hypofunction) with also abnormalities in thalamus and basal ganglia reported. | Kaminski *et al.*(110); Lee and Park, (111); Manoach *et al.* (112); Schlösser *et al.* (113); Schneider *et al.* (114); Borgan *et al.* (115); Eryilmaz *et al.* (116) |
| ***Long-term memory and language*** | | | | | | |
| Semantic memory (SM) | The long term storage of personally relevant semantic knowledge, independent of recalling a specific experience | Activated during SM contrasts: experiments mainly comprising paradigms: words vs. pseudo words, semantic vs. phonological task, high vs. low meaningfulness | 23 | Binder, (117) | Semantic memory-based processing is impaired in SCZ, e.g., semantic retrieval, encoding and association, and were found to associate with deficits (e.g., increased connectivity and decreased neural activation) in fronto-parieto-temporal network (e.g., inferior parietal lobule, medial/inferior prefrontal gyrus, and superior/middle temporal gyrus) and relate to negative and positive symptoms and formal thought disorder. | Jamadar *et al.* (118,119) Kubicki *et al.* (120) Ragland *et al.* (121) |
| Speech production (SP) | The process by which thoughts are translated into speech, involving the integration of auditory, somatosensory, and motor information | Studies contrasted speech production (including phonemes, syllables, words, sentences or narratives) with a condition in which no speech was produced | 13 | Adank, (122) | Abnormal speech production in schizophrenia contributes to the symptom of formal thought disorder (FTD). FTD-associated production of disorganized speeches was correlated with activity in fusiform, inferior frontal and superior temporal cortex. Reversed laterality of activation in the lateral temporal cortex was found in schizophrenia patients during speech production, which has been related to glutamatergic imbalance. | Kircher *et al.* (123) Nagels *et al.* (124) McGuire *et al.* (125) |

| | | | | | | |
|---|---|---|---|---|---|---|
| Autobiographic Memory (AM) | Long-term memory for personal experiences and personal knowledge of an individual's life | Tasks referring to autobiographical recall: episodic recollection of personal events from one's own life | 23 | Spreng, (126) | AM is impaired (e.g., reduced specificity and retrieval of memories) in schizophrenia and has been found to associate with reduced hippocampal volume and altered neural activations in multiple brain regions (e.g., anterior cingulate cortex, and lateral prefrontal cortex). | Herold *et al.* (127)<br>Cuervo-Lombard *et al.* (128)<br>Herold *et al.* (129) |
| ***Sensory-motor*** | | | | | | |
| Motor | Motor execution | Finger tapping > baseline; excl. regions associated with visually paced finger-tapping tasks | 10 | Witt *et al.* (130) | Motor cortex and its closely inter-connected brain regions (e.g., prefrontal–motor, cerebello-thalamo-motor, sensory-motor and basal ganglia circuits) showed abnormalities in schizophrenia, and were related to motor behavioral problems observed in the patients including motor learning, sequential movements and postural control, and have also been associated with clinical symptoms. | Walther *et al.* (131)<br>Marvel *et al.* (132)<br>Berman *et al.* (133)<br>Bernard *et al.* (134)<br>Du *et al.* (135) |
| Auditory | Auditory sensory processing | Purely auditory tasks using highly controlled synthesized acoustic stimuli | 11 | Petacchi *et al.* (136) | Auditory processing including automatic, feed forward and pre-attentive functions are impaired in schizophrenia. The auditory oddball tasks revealed multiple regions within the auditory network that were abnormally activated, e.g., the (middle/superior) temporal cortex, insula, prefrontal and inferior parietal cortex, and the abnormalities were associated with negative symptoms and cognitive deficits. | Sweet *et al.* (137)<br>Perez *et al.* (138)<br>Force *et al.* (139)<br>Shin *et al.* (140)<br>Wolf *et al.* (141)<br>Kim *et al.* (142)<br>Shim *et al*. (143) |

[a]Details on the Identification of DMN regions involved in socio-affective processing can be found at (144); [b]the MDN network was derived from a conjunction across three neuroimaging meta-analyses on working memory, vigilant attention, and inhibitory control using coordinate-based ALE (145). ALE: activation likelihood estimation; MACM: meta-analytic connectivity modeling.

**Table S4. Functional MRI scanning parameters for each site**

| Site | Scanner Type | Magnetic Field | TR (ms) | TE (ms) | FA (°) | No. Slices | Voxel-size ($mm^3$) | Orientation | Scan Duration (s) |
|---|---|---|---|---|---|---|---|---|---|
| ***The main, international cohort*** | | | | | | | | | |
| Aachen-1 | Siemens TrioTim | 3.0T | 2000 | 28 | 77 | 34 | 3.3 3.6 x 3.6 | Axial | 412 |
| Aachen-2 | Siemens TrioTim | 3.0T | 2000 | 21 | n.a. | 44 | 3 x 3 x 3 | Axial | 480 |
| Albuquerque (COBRE) | Siemens TrioTim | 3.0T | 2000 | 29 | 75 | 32 | 4 x 3 x 3 | Axial | 300 |
| Göttingen | Siemens TrioTim | 3.0T | 2000 | 30 | 70 | 33 | 3 x 3 x 3 | Axial | 312 |
| Groningen | Philips Achieva | 3.0T | 2400 | 28 | 85 | 43 | 3 x 3.44 x 3.44 | Axial | 480 |
| Utrecht | Philips Achieva[1] | 3.0T | 21.75[1] | 32.4 | 10 | 40 | 4 x 4 x 4 | Coronal | 363 |
| Lille | Philips[1] Achieva | 3.0T | 19.25 | 9.6 | 9 | 45 | 3.22 x 3.22 x 3.4 | Sagittal | 896 |
| ***The validation B-SNIP sample*** | | | | | | | | | |
| Hartford | Siemens Allegra | 3.0T | 1500 | 27 | 70 | 29 | 3.4x3.4x5 | Axial | 315 |
| Baltimore | Siemens Triotim | 3.0T | 2210 | 30 | 70 | 36 | 3.4x3.4x3 | Axial | 309.4 |

*Note:* TR: repetition time, TE: echo time, FA: flip angle; [1]PRESTO-SENSE sequence combining a 3D-PRESTO pulse sequence with parallel imaging in 2 directions (8-channel SENSE head-coil) which achieved full brain coverage within 609 ms for the Utrecht site and within 1001 ms for the Lille site.

**Table S5. T1-weighted structural MRI scanning parameters for each site**

| Site | Scanner Type | Magnetic Field | TR (ms) | TE (ms) | FA (°) | No. Slices | Voxel-size ($mm^3$) |
|---|---|---|---|---|---|---|---|
| ***The main, international cohort*** | | | | | | | |
| Aachen-1 | Siemens TrioTim | 3.0T | 2300 | 3.03 | 9 | 176 | 1 x 1x 1 |
| Aachen-2 | Siemens TrioTim | 3.0T | 1900 | 2 | n.a. | 176 | 0.97 x 0.97x1 |
| Albuquerque (COBRE) | Siemens TrioTim[1] | 3.0T | 2530 | [1.64, 3.5, 5.36, 7.22, 9.08] | 7 | 176 | 1 x 1x 1 |
| Göttingen | Siemens TrioTim | 3.0T | 2250 | 3.26 | n.a. | 176 | 1 x 1x 1 |
| Groningen | Philips Achieva | 3.0T | 2500 | 4.6 | 30 | 160 | 1 x 1x 1 |
| Utrecht | Philips Achieva[2] | 3.0T | 9.86 | 4.6 | n.a. | 160 | 0.875 x 0.875 x 1 |
| Lille | Philips Achieva[2] | 3.0T | 10 | 4.6 | n.a. | 160 | 1 x 1x 1 |
| ***The validation B-SNIP sample*** | | | | | | | |
| Hartford | Siemens Allegra | 3.0T | 2300 | 2.91 | 9 | 160 | 1 x 1x1.2 |
| Baltimore | Siemens Triotim | 3.0T | 2300 | 2.91 | 9 | 160 | 1 x 1x1.2 |

*Note:* TR: repetition time, TE: echo time, FA: flip angle; [1]a multi-echo MPRAGE (MEMPR) sequence with 5 TEs; [2]PRESTO-SENSE sequence.

**Table S6. The identified reliably relevant connections for the ToM and the eSAD networks in the prediction of the cognitive dimension**

| Network | Connection |
|---|---|
| **ToM** | vmPFC<->PCC/PrC; vmPFC<->right pSTS; PCC/PrC<->left MTG; TPJ<->dmPFC; left MTG<-> left aMTG; left MTG<->RIFG; RMTG<-> left lFG; right MTG<-> right aMTG |
| **eSAD** | ACC<-> right Amy; SGC<->PCC/PrC; SGC<->dmPFC; PCC<-> left aMTG; PCC<->vmPFC; dmPFC<->rvBG; dmPFC<-> left Amy; vmPFC<->RTPJ; left vBG<-> left Amy; right Amy<-> left Amy |

Abbreviations: ToM, theory-of-mind; eSAD, extended socio-affective default. Amy, amygdala; Hipp, hippocampus; vmPFC, ventro-medial prefrontal cortex; dmPFC, dorso-medial prefrontal cortex; dmPFG, dorso-medial prefrontal cortex; aMTG, anterior middle temporal gyrus, IFG, inferior frontal gyrus; TPJ, temporo-parietal junction, PCC, posterior cingulated cortex, PrC, precuneus; SGC, subgenual cingulate cortex, vBG, ventral basal ganglia; ACC, anterior cingulated cortex.

**Table S7. Coordinates and brain locations of the nodes connected by reliably predictive connections within the identified robustly predictive networks**

| Network | MNI coordinates x | y | z | Macroanatomy of nodes | Macroanatomy of the connected nodes |
|---|---|---|---|---|---|
| ***ToM*** | | | | | |
| | **0** | **52** | **-12** | **vmPFC** | PrC; right pSTS |
| | **2** | **-56** | **30** | **PCC/PrC** | vmPFC; left MTG |
| | 50 | -34 | 0 | pSTS | vmPFC |
| | 56 | -50 | 18 | TPJ | dmPFC |
| | -8 | 56 | 30 | dmPFC | Right TPJ |
| | -54 | -28 | -4 | MTG | PrC; left aMTG; right IFG |
| | 52 | -18 | -12 | MTG | left IFG; right aMTG |
| | 54 | -2 | -20 | aMTG | right MTG |
| | -54 | -2 | -24 | aMTG | left MTG |
| | -48 | 30 | -12 | IFG | right MTG |
| | 54 | 28 | 6 | IFG | left MTG |
| ***eSAD*** | | | | | |
| | 0 | 38 | 10 | ACC | right Amy/Hipp |
| | -2 | 32 | -8 | SGC | PCC/PrC; dmPFC |
| | -2 | -52 | 26 | PCC/PrC | SGC; left aMTG; vmPFC |
| | -2 | 52 | 14 | dmPFC | SGC; right vBG; left Amy/Hipp |
| | -54 | -10 | -20 | aMTG | PCC/PrC |
| | -2 | 50 | -10 | vmPFC | PCC/PrC; right TPJ |
| | 6 | 10 | -8 | vBG | dmPFC |
| | -6 | 10 | -8 | vBG | left Amy/Hipp |
| | -24 | -10 | -20 | Amy/Hipp | left vBG; dmPFC; right Amy/Hipp |
| | 24 | -8 | -22 | Amy/Hipp | left Amy/Hipp |
| | 50 | -60 | 18 | TPJ | vmPFC |

*Note:* Coordinate (x, y, z) of each node is reported in standard space of the Montreal Neurological Institute (MNI) as demonstrated in the source publications of the two identified functional networks. Nodes that were spatially overlapping between the subnetworks of ToM and eSAD are highlighted in red.

Abbreviations: ToM, theory-of-mind; eSAD, extended socio-affective default. Amy, amygdala; Hipp, hippocampus; vmPFC, ventro-medial prefrontal cortex; dmPFC, dorso-medial prefrontal cortex; dmPFG, dorso-medial prefrontal cortex;  aMTG, anterior middle temporal gyrus, IFG, inferior frontal gyrus; TPJ, temporo-parietal junction, PCC, posterior cingulated cortex, PrC, precuneus; SGC, subgenual cingulate cortex, vBG, ventral basal ganglia; ACC, anterior cingulated cortex.

**Table S8.** **Node importance for the ToM and the eSAD networks**

| Network | Node | Importance score |
|---|---|---|
| **ToM** | vmPFC | 4.20 |
| | mFG | 3.39 |
| | dmPFC | 3.33 |
| | PCC/PrC | 4.24 |
| | Right TPJ | 2.83 |
| | Left TPJ | 3.67 |
| | Right aMTG | 3.89 |
| | Left aMTG | 4.81 |
| | rMTG | 4.58 |
| | lMTG | 4.87 |
| | Right pSTS | 3.11 |
| | lpSTS | 2.82 |
| | rIFG | 3.07 |
| | lIFG | 4.07 |
| | rV5 | 3.15 |
| **eSAD** | ACC | 3.21 |
| | SGC | 4.14 |
| | PCC | 4.59 |
| | dmPFC | 4.05 |
| | rTPJ | 1.96 |
| | lTPJ | 2.56 |
| | lvBG | 4.40 |
| | rvBG | 3.50 |
| | Left aMTG | 2.85 |
| | Right Amy | 4.27 |
| | Left Amy | 3.98 |
| | vmPFC | 4.21 |

Abbreviations: ToM, theory-of-mind; eSAD, extended socio-affective default. Amy, amygdala; Hipp, hippocampus; vmPFC, ventro-medial prefrontal cortex; dmPFC, dorso-medial prefrontal cortex; dmPFG, dorso-medial prefrontal cortex; aMTG, anterior middle temporal gyrus, IFG, inferior frontal gyrus; TPJ, temporo-parietal junction, PCC, posterior cingulated cortex, PrC, precuneus; SGC, subgenual cingulate cortex, vBG, ventral basal ganglia; ACC, anterior cingulated cortex.

**Figure S1. Illustration of the robust four-dimensional representation of psychopathology identified in our prior machine-learning study (A) and the dimensional-scores estimated for the main (B) and the validation (C) samples investigated in the present study**

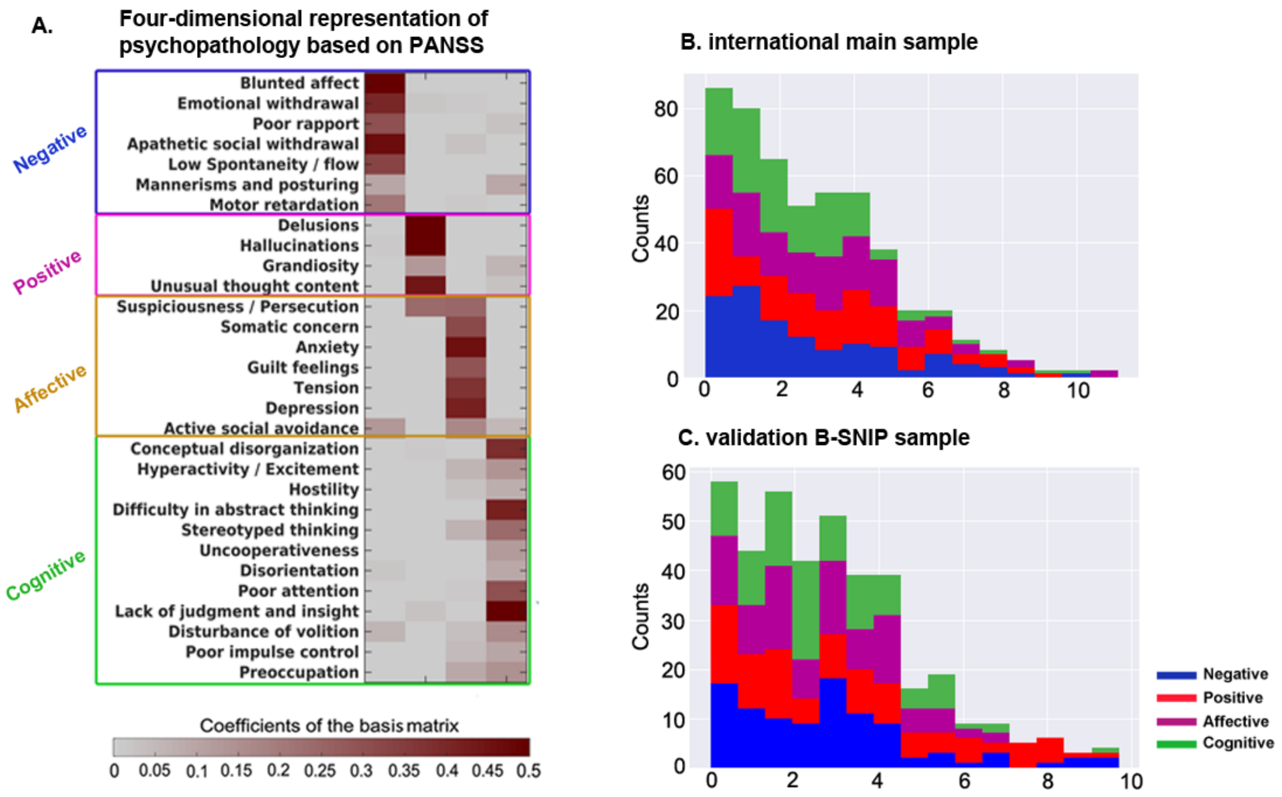

The four dimension model shown in the left panel was adapted from Figure 1 in (15) with permission. The sample used for creating the four dimension model is independent of the main and the validation samples analyzed in the present work. PANSS: Positive and Negative Syndrome Scale.

**Figure S2. Multivariable prediction of the original three PANSS subscales from the resting-state functional connectivity within each of the 17 functional networks using the same validation procedure as we have done for the four symptom dimensions**

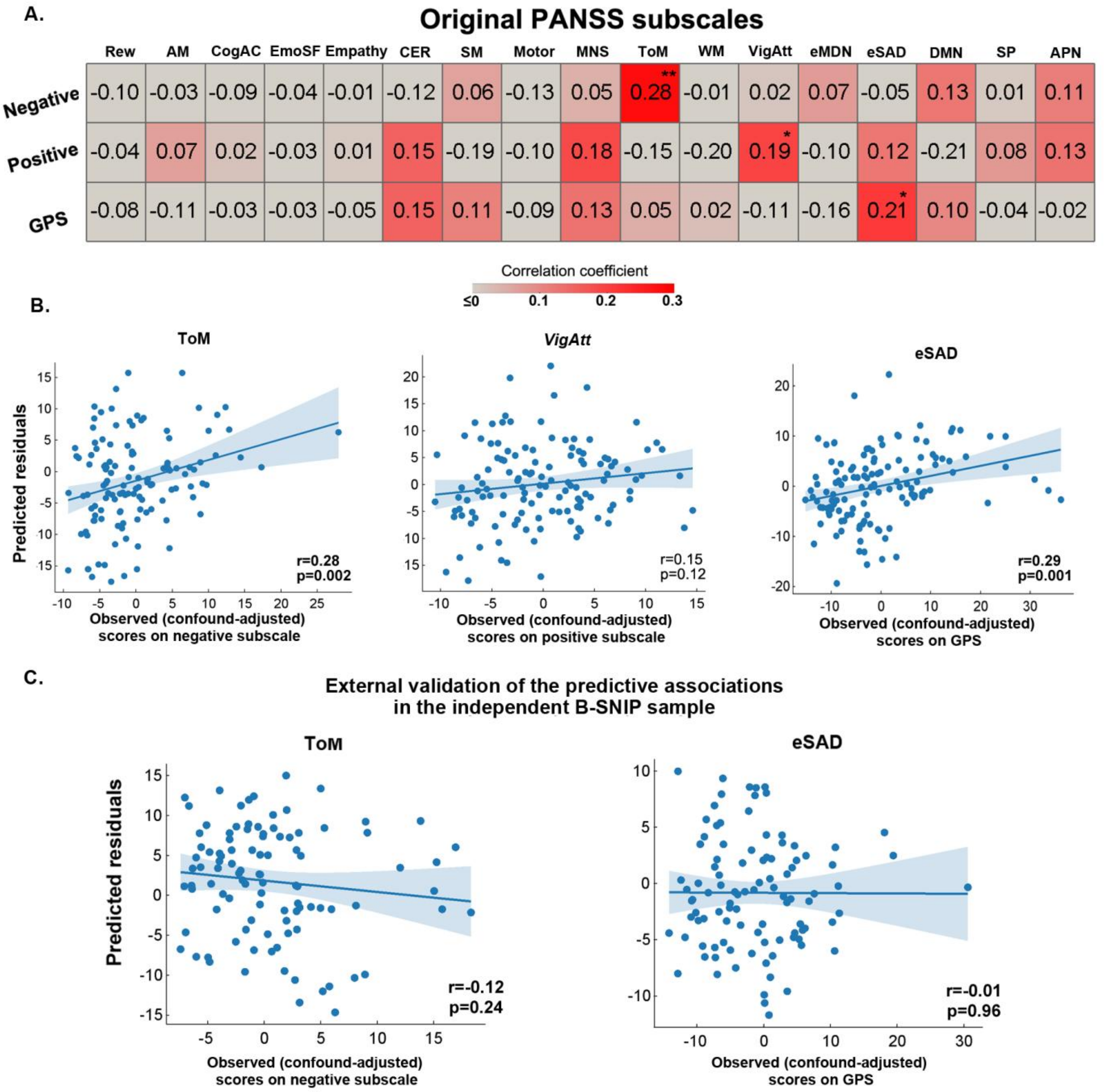

**A)** Tile plot shows the 10-fold cross-validation results for the main sample in the prediction of the three PANSS subscales. **p*<0.05, ***p*<0.01, identified through 1000 permutation tests.

**B)** Scatter plots show the leave-one-site-out cross-validation results for the three significant predictions identified in 500x repeated 10-fold cross-validation in the main sample. Except for the prediction of the positive subscale from the rs-FC within the vigilant attention network, other two predictions were both confirmed by leave-one-site-out cross-validation with significant correlations observed.

**C)** Scatter plot show that neither of the two tested predictive patterns was significant in the B-SNIP sample.

Abbreviations: EmoSF, emotional scene and face processing; Rew, reward-related decision making; CER, cognitive emotion regulation; ToM, theory-of-mind; MNS, minor neuron system; DMN, default mode network; eSAD, extended socio-affective default; VigAtt, vigilant attention; CogAC, cognitive action control; eMDN, the extended multi-demand networks; SM, semantic memory; SP, speech production; WM, working memory; AM, autobiographical memory; APN, auditory processing network. GPS: general psychopathology subscale.

**Figure S3.** **Histograms of the three metrics assessing the property of random networks**

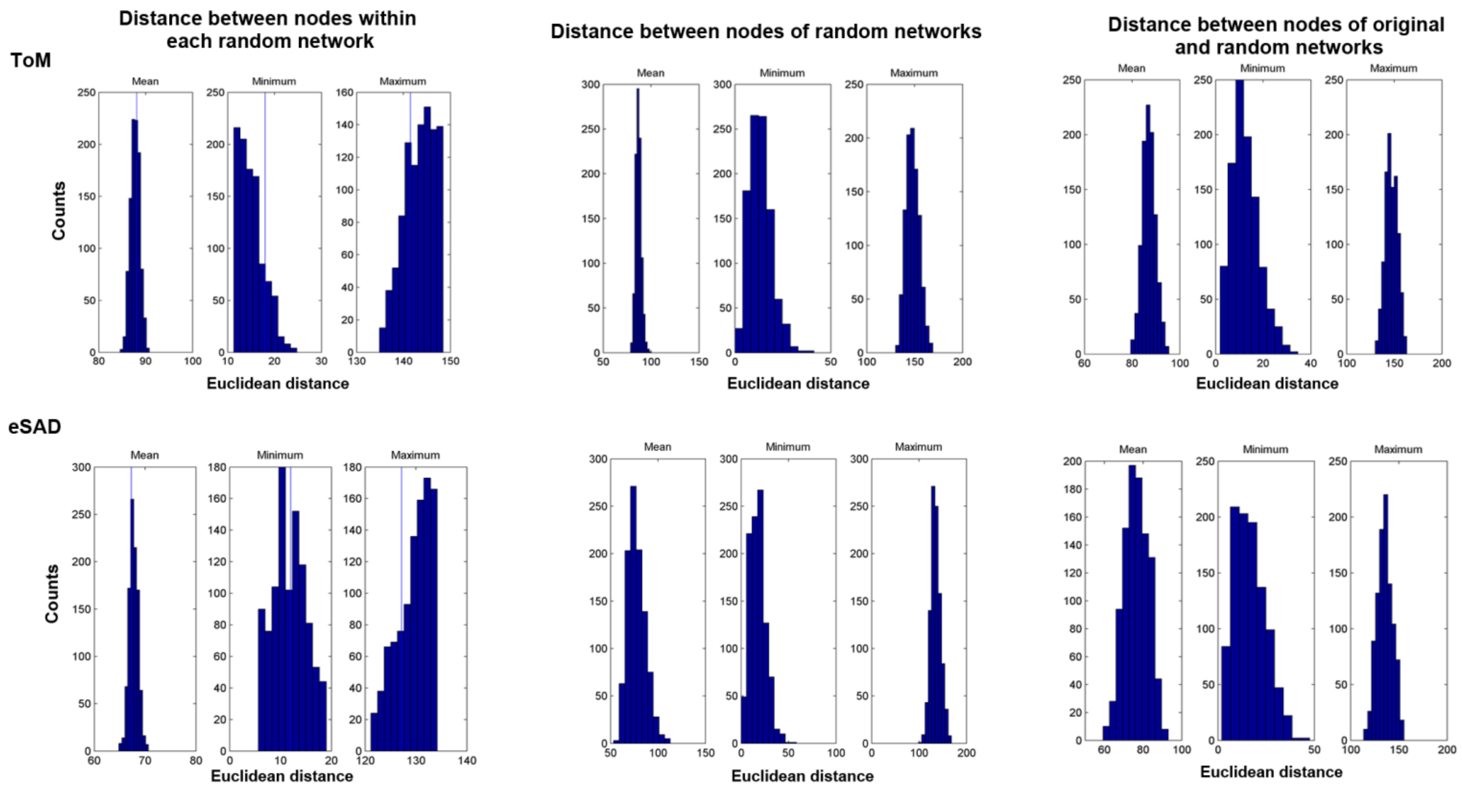

Abbreviations: ToM, theory-of-mind; eSAD, extended socio-affective default

**Figure S4. Reliably predictive connections for the theory-of-mind (ToM) network in the prediction of the negative dimension and the two subnetworks within ToM**

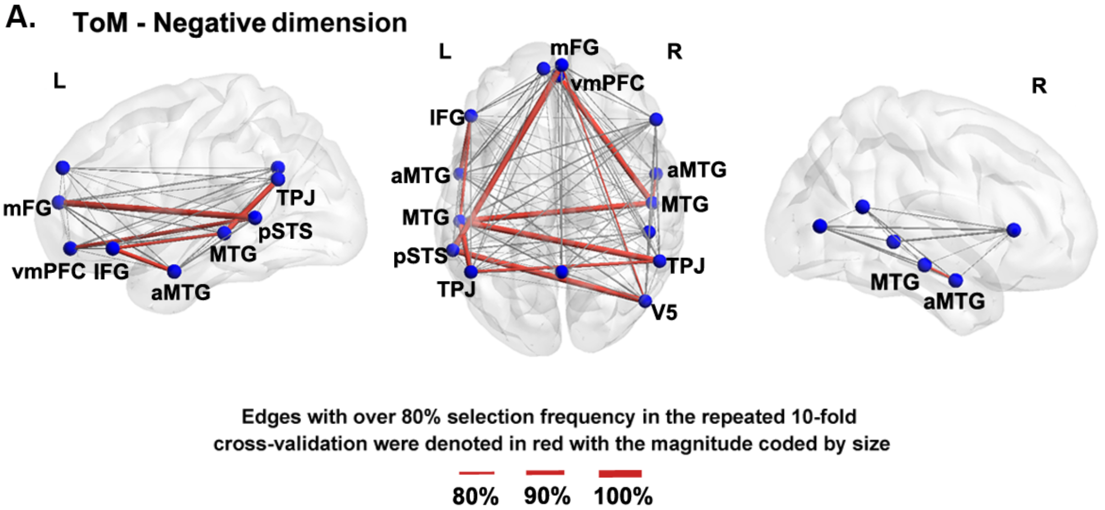

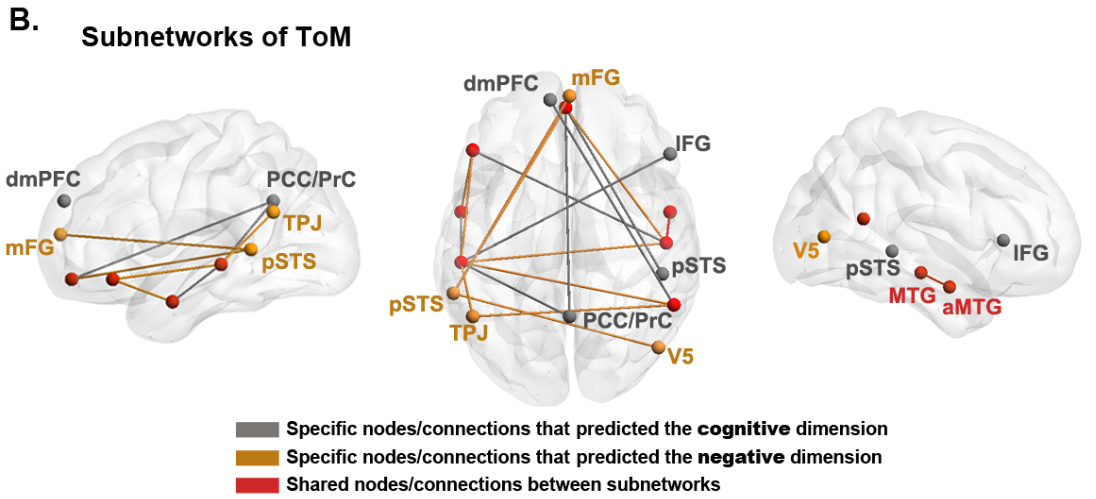

**A)** Reliably relevant connections selected by 10-fold process on main sample in the prediction of negative dimension. The reliably relevant edges are colored in red and the selection frequency for these edges was coded by line size. Other connections within each of the networks are shown in light grey. These connections were all reliably selected in both the seven leave-one-site-out experiments and the models trained within the entire main sample for validation in B-SNIP.

**B)** Subnetworks of ToM which predicted the cognitive or the negative dimension of psychopathology. Their shared nodes and connections were shown in red color.

correlation and regression analyses. *Behav Res Methods* 41(4): 1149-1160.

14. Power JD, Barnes KA, SnyderAZ, Schlaggar BL, Petersen SE (2012): Spurious but systematic correlations in functional connectivity MRI networks arise from subject motion. *Neuroimage* 59: 2142-2154.
15. Chen J, Patil K R, Weis S, Sim K, Nickl-Jockschat T, Zhou J, *et al.* (2020): Neurobiological Divergence of the Positive and Negative Schizophrenia Subtypes Identified on a New Factor Structure of Psychopathology Using Non-negative Factorization: An International Machine Learning Study. *Biol Psychiatry* 87(3): 282-293
16. Power JD, Schlaggar BL, Petersen SE (2015): Recent progress and outstanding issues in motion correction in resting state fMRI. *Neuroimage* 105: 536-551.
17. Parker D, Liu X, Razlighi QR (2017): Optimal slice timing correction and its interaction with fMRI parameters and artifacts. *Med Image Anal* 35: 434-445
18. Calhoun VD, Wager TD, Krishnan A, Rosch KS, Seymour KE, Nebel MB, *et al.* (2017): The impact of T1 versus EPI spatial normalization templates for fMRI data analyses. *Hum Brain Mapp* 38: 5331-5342.
19. Varikuti DP, Hoffstaedter F, Genon S, Schwender H, Reid AT, Eickhoff SB (2017): Resting-state test–retest reliability of a priori defined canonical networks over different preprocessing steps. *Brain Struct Funct* 222: 1447-1468.
20. Murphy K, Fox MD (2017): Towards a consensus regarding global signal regression for resting state functional connectivity MRI. *Neuroimage* 154: 169-173.
21. Liu TT, Nalci A, Falahpour M (2017): The global signal in fMRI: Nuisance or Information? *NeuroImage* 150: 213-229.
22. Yang G J, Murray J D, Repovs G, Cole MW, Savic A, Glasser MF, *et al.* (2014): Altered global brain signal in schizophrenia. *Proc Natl Acad Sci U S A* 111(20): 7438-7443.
23. Tipping ME (2001): Sparse Bayesian learning and the relevance vector machine. *J Machine Learn Res* 1: 211-244.
24. Soentpiet R (1999): Advances in kernel methods: support vector learning. MIT press.
25. Ghosh S, Mujumdar PP (2008): Statistical downscaling of GCM simulations to stream flow using relevance vector machine. *Adv Water Resour* 31: 132-146.
26. Zheng Y-T, Neo S-Y, Chua T-S, Tian Q (2018): Probabilistic optimized ranking for multimedia semantic concept detection via RVM. *Proceedings of the 2008 international conference on Content-based image and video retrieval*. Niagara Falls, Canada: ACM. pp. 161–168.

27. Sripada C, Rutherford S, Angstadt M, Thompson WK, Luciana M, Weigard A, *et al.* (2019): Prediction of neurocognition in youth from resting state fMRI. *Mol Psychiatry* doi: 10.1038/s41380-019-0481-6.

28. More S, Eickhoff SB, Julian C, Patil, KR (2020): Confound Removal and Normalization in Practice: A Neuroimaging Based Sex Prediction Case Study. Preprint available at https://juser.fz-juelich.de/record/877721; Accepted in the *European Conference on Machine Learning and Principles and Practice of Knowledge Discovery in Databases* (ECML PKDD; https://ecmlpkdd2020.net/programme/accepted/#ADSTab).

29. Drysdale AT, Grosenick L, Downar J, Dunlop K, Mansouri F, Meng Y, *et al.* Resting-state connectivity biomarkers define neurophysiological subtypes of depression. *Nat Med* 23(1): 28-38.

contribution of the GABA-benzodiazepine α1 receptor subtype to [11C] Ro15-4513 PET images. *J Cereb Blood Flow Metab* 32: 731–744.

62. Kim M, Kang BN, Lim JY (2016): Decision-making deficits in patients with chronic schizophrenia: Iowa

Gambling Task and Prospect Valence Learning model. *Neuropsychiatr Dis Treat* 12: 1019–1027.

88. Park S, Gibson C, McMichael T (2006): Socioaffective factors modulate working memory in schizophrenia

patients. *Neuroscience* 139: 373-384.

89. Laird AR, Eickhoff SB, Li K, Robin DA, Glahn DC, Fox PT (2009): Investigating the functional heterogeneity of the default mode network using coordinate-based meta-analytic modeling. *J Neurosci* 29(46): 14496-14505.
90. Garrity AG, Pearlson GD, McKiernan K, Lloyd D, Kiehl KA, Calhoun VD (2007): Aberrant “default mode” functional connectivity in schizophrenia. *Am J Psychiatry* 164(3): 450-457.
91. Hu ML, Zong XF, Mann JJ, Zheng J, Liao Y, Li Z, *et al.* (2017): A review of the functional and anatomical default mode network in schizophrenia. *Neurosci Bull* 33(1): 73-84.
92. Jia W, Zhu H, Ni Y, Su J, Xu R, Jia H, Wan X Disruptions of frontoparietal control network and default mode network linking the metacognitive deficits with clinical symptoms in schizophrenia. *Hum Brain Mapp* doi: 10.1002/hbm.24887.
93. Pomarol-Clotet E, Salvador R, Sarro S, Gomar J, Vila F, Martinez A, *et al.* (2008): Failure to deactivate in the prefrontal cortex in schizophrenia: dysfunction of the default mode network?. *Psychol Med* 38(8): 1185-1193.
94. Du Y, Pearlson GD, Yu Q, He H, Lin D, Sui J, *et al.* (2016): Interaction among subsystems within default mode network diminished in schizophrenia patients: a dynamic connectivity approach. *Schizophr Res* 170(1): 55-65.
95. Langner R, Eickhoff SB (2013): Sustaining attention to simple tasks: a meta-analytic review of the neural mechanisms of vigilant attention. *Psychol Bull* 139(4): 870-900.
96. Eyler LT, Olsen RK, Jeste DV, Brown GG (2004): Abnormal brain response of chronic schizophrenia patients despite normal performance during a visual vigilance task. *Psychiatry Res Neuroimaging* 130(3): 245-257.
97. O’Gráda C, Barry S, McGlade N, Behan C, Haq F, Hayden J, *et al.* (2009): Does the ability to sustain attention underlie symptom severity in schizophrenia? *Schizophr Res* 107(2-3): 319-323.
98. Cieslik EC, Mueller VI, Eickhoff CR, Langner R, Eickhoff SB (2015): Three key regions for supervisory attentional control: evidence from neuroimaging meta-analyses. *Neurosci Biobehav Rev* 48: 22-34.
99. Reuter B, Jäger M, Bottlender R, Kathmann N (2007): Impaired action control in schizophrenia: the role of volitional saccade initiation. *Neuropsychologia* 45(8): 1840-1848.
100. Braver TS, Barch DM, Cohen JD (1999): Cognition and control in schizophrenia: a computational model of dopamine and prefrontal function. *Biol Psychiatry* 46(3): 312-328.
101. Barch D M, Sheffield JM (2017): *Cognitive control in schizophrenia: Psychological and neural mechanisms*, In T. Egner (Ed.), *The Wiley handbook of cognitive control.* Wiley-Blackwell, pp 556–580.
102. Minzenberg MJ, Lesh TA, Niendam TA, Yoon JH, Rhoades RN, Carter CS (2014): Frontal cortex control dysfunction related to long-term suicide risk in recent-onset schizophrenia. *Schizophr Res* 157(1-3): 19-25.

109. Rottschy C, Langner R, Dogan I, Reetz K, Laird AR, Schulz JB, *et al.* Modelling neural correlates of working memory: a coordinate-based meta-analysis. *NeuroImage* 60(1): 830-846.

10.1038/s41380-019-0619-6.

autobiographical memory, and hippocampal volume in “younger” and “older” patients with chronic schizophrenia. *Front Psychiatry* 6: 53
